# Cognitive function depends upon *Satb2* gene dosage in cortical projection neurons

**DOI:** 10.64898/2026.06.13.731750

**Authors:** Thomas S. Finn, Jeremiah Tsyporin, Min Dai, Sherry Jingjing Wu, Gabrielle Servito, Xiaokuang Ma, John Zhang, Esther Park, Charlene Guo, Austin Schubert, Hailey Lizarraga, Ariella Angelini Stewart, Vipin Kumar, Giovanni J. Marrero, Yufei Si, Sol Katzman, Alejandro Fillon, Sofia Martin, Jiang Wu, Yuri Zarate, Fei Chen, Evan Macosko, Lu Chen, Shenfeng Qiu, Gord Fishell, Bin Chen

## Abstract

SATB2-associated syndrome (SAS) is a severe neurodevelopmental disorder caused by *de novo* heterozygous *SATB2* mutations, yet how haploinsufficiency disrupts brain development remains poorly understood. While homozygous *Satb2* loss causes profound embryonic cell-fate defects, we demonstrate using a heterozygous mouse model that SAS phenotypes emerge primarily during postnatal circuit maturation. Integrating chromatin profiling, transcriptomics, electrophysiology, and behavior, we show that SATB2 acts as a dose-sensitive chromatin regulator that binds conserved enhancer–promoter landscapes to orchestrate networks linked to human intelligence. Although excitatory neuron subtype specification is preserved, *Satb2* heterozygotes adopt an intermediate epigenetic state that drives cell-type-specific dysregulation of genes enriched for intellectual disability risk variants. Consequently, mutant neurons exhibit simplified dendritic arborization, reduced intrinsic excitability, and weakened layer 2/3-to-layer 5 intracortical connectivity. These circuit deficits culminate in the disorganization of the somatosensory barrel cortex and severe impairments in whisker-dependent texture discrimination. Finally, by restricting *Satb2* heterozygosity to the cortex, we decouple these cortical sensory deficits from subcortical vocalization phenotypes. Together, our work links SATB2 dosage to chromatin architecture and postnatal circuit maturation, revealing a critical, post-mitotic therapeutic window for intervention in SAS.

## Introduction

SATB2-Associated Syndrome (SAS) is a neurodevelopmental disorder characterized by craniofacial defects, behavioral abnormalities, and profound intellectual disability with absent or limited speech^1,2^. SAS is caused by *de novo* heterozygous mutations in *SATB2*, a DNA-binding protein that organizes higher-order chromatin structure and recruits chromatin remodeling complexes to regulate transcription^3^. Because brain expression of SATB2 is largely restricted to excitatory neurons (ExNs) in the cerebral cortex and hippocampus^4^, the cognitive deficits of SAS points to a critical, dose-dependent requirement for SATB2 during cortical development.

The six-layered neocortex integrates sensory information to drive complex cognitive behaviors^5^. Within these circuits, ExNs serve as the primary mediators of input/output signaling and are broadly categorized by their axon targets: layer 6 corticothalamic (CT) neurons, layer 5b subcerebral pyramidal tract (PT) neurons, and intra-telencephalic (IT) neurons distributed across layers 2-6^6,7^. While single-cell transcriptomics has precisely refined these subtype classifications^8,9^, our mechanistic understanding of how their distinct identities are specified relies almost exclusively on homozygous knockout mouse models. In the embryonic cortex, SATB2 specifies the cortical callosal– and subcerebral-projecting neurons^10–12^ through its actions in gene activation and repression^12^. By recruiting the nucleosome remodeling and deacetylase (NuRD) complex, SATB2 represses the expression of *Bcl11b* in callosal neurons, a gene essential for axonal projection of L5 PTs^10,11^. SATB2 is also required for the expression of *Fezf2*, a gene encoding a transcription factor necessary for specifying L5 PT identity^13–17^. Consequently, homozygous loss of *Satb2* leads to the absence of L5 PTs^12^.

Over 120 unique *SATB2* variants have been identified in SAS patients, with approximately 70% representing likely loss-of-function (LOF) alleles^1,2,18^. However, the developmental mechanisms disrupted by human *SATB2* haploinsufficiency remain unknown. Because of this complete lack of mechanistic clarity, current therapeutic options for SAS are strictly symptom-guided, focusing primarily on pediatric dentistry and speech therapy^18^. Defining the dose-dependent gene regulatory networks governed by SATB2 is therefore essential to uncovering fundamental tenets of cortical assembly and identifying actionable therapeutic targets.

Here, we leverage chromatin profiling, single-nucleus transcriptomics, and spatial transcriptomics in a mouse model of SAS to reveal how SATB2 dosage controls cortical maturation. We show that SATB2 binds the promoters and enhancers of genes linked to human intelligence across all ExN subtypes. Beyond the NuRD complex, we demonstrate that SATB2 interacts with the mSWI/SNF (BAF) chromatin-remodeling complex and cooperates with key cortical transcription factors, including CUX1, BCL11A, and LHX2, to execute its dual roles in activation and repression. Crucially, *Satb2* haploinsufficiency does not disrupt embryonic cell-fate specification as seen in null mutants; instead, it establishes an intermediate epigenetic state that postnatally dysregulates intelligence-associated gene networks. These molecular shifts culminate in persistent deficits in dendritic arborization, altered neuronal excitability, defective intracortical circuitry, and impaired sensory behavior. Together, our findings map the dose-sensitive architecture of cortical development and illuminate a postnatal therapeutic window for SAS.

## Results

### SATB2 binds to promoters and enhancers of genes associated with intelligence

We identified transcriptional targets of SATB2 by performing chromatin immunoprecipitation followed by high-throughput DNA sequencing (ChIP-seq) using P0 wildtype cortices and SATB2 antibodies (**Extended Data Fig. 1a-b**). We also performed Cleavage Under Targets & Release Using Nuclease (CUT&RUN)^19,20^ experiments using antibodies for SATB2. After mapping sequencing reads to the genome, binding peaks were called using MACS2^21^ (for ChIP-seq) and SEACR^19^ (for CUT&RUN). We identified SATB2 binding peaks at both promoters (defined as less than 3kb upstream and downstream from the transcription start site, TSS) and potential enhancer regions (defined as greater than 3kb upstream and downstream from the TSS) (**Extended Data Fig. 1a**). To quantify the concordance between SATB2 ChIP-seq and CUT&RUN datasets, we stratified the binding sites into promoter and distal intergenic enhancers using ChIP-seeker. We then performed a permutation test (*n* = 1,000 iterations) using the regioneR package to assess whether the observed overlaps significantly exceeded those by random chance. This analysis revealed a significant concordance in both genomic contexts. Specifically, promoter regions yielded a Z-score of 535.6 (*p* < 0.001), while distal enhancers yielded a Z-score of 476.0 (*p* < 0.001). The magnitude of these Z-scores confirms that the SATB2 binding profiles derived from CUT&RUN are consistent with ChIP-seq data (**Extended Data Fig 1b**). To visually illustrate ChIP-seq and CUT&RUN reproducibility, we selected genes that have strong SATB2 binding peaks and aligned the respective tacks in the IGV browser (**Extended Data Fig 1c**).

To characterize cell-type specific SATB2 binding, we integrated Satb2 ChIP-seq and CUT&RUN peaks with publicly available E18 mouse brain single nuclei multiome (Gene Expression+ATAC in the same cell; snMultiome) sequencing data^22^. We observed enrichment of SATB2 binding in accessible regions in upper-, and deep-layer (TCERG1L, L5 PT and HS3ST4, L6 CT) ExNs (**Extended Data Fig. 2a).** To determine the disease-relevance of the SATB2 genomic targets, we first applied cell type-stratified linkage disequilibrium score regression (LDSC)^23^ to the E18 mouse brain snMultiome sequencing data to quantify the enrichment of phenotype heritability in different cell types based on summary statistics from multiple genome-wide association studies (GWAS)^24^. This analysis revealed numerous disease-associated risk variants enriched in the accessible regions in specific cell-types, notably for variants associated with intelligence and schizophrenia in upper-layer and deep-layer ExNs (**Fig. 1a**). The SATB2 binding regions were enriched in variants associated with neurodevelopmental disorders, especially in those associated with intelligence, schizophrenia, and bipolar disorder (**Fig. 1a**). Interestingly, enrichment is stronger in the enhancer regions than the promoter regions (**Fig. 1a**). These findings suggest that SATB2 extensively regulates promoters and enhancers of genes associated with intelligence and cognitive disorders.

**Figure 1:**
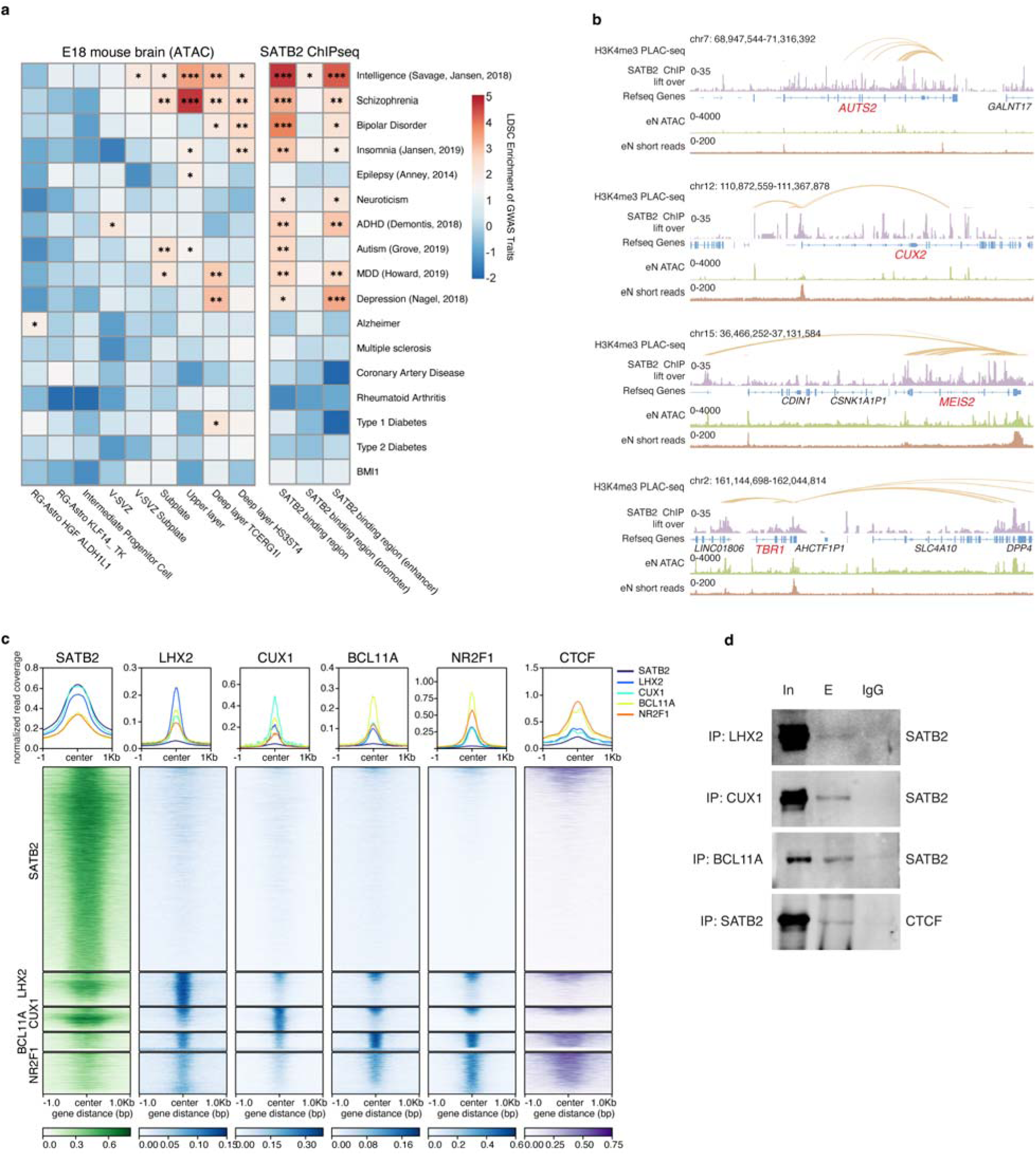
SATB2 binds to promoters and/or enhancers of genes associated with human neurodevelopmental diseases and interacts with other cortical transcription factors. **a**, Cell type-stratified lineage disequilibrium score regression (LDSC) heatmaps based on summary statistics from GWAS for snMultiome data and SATB2 ChIP-seq peaks.\**p* < 0.05; \*\**p* < 0.01; \*\*\**p* < 0.001. Exact *p* values are provided in **Supplementary Table 2**. **b**, Human genome tracks (hg38) showing SATB2 binds to the promoters and enhancers of ID genes (shown in red). The mouse SATB2 ChIP-seq data liftover, human H3K4me3-PLAC-seq, ATAC-seq and RNA-seq for enriched human fetal excitatory populations are shown^26^. ID genes *AUTS2*, *CUX2*, *MEIS2*, and *TBR1* are shown as examples**. c**, Top, normalized read coverage showing SATB2, LHX2, CUX1, BCL11A, and NR2F1 CUT&RUN peaks overlapping with each other, and the CTCF ChIP-seq peaks overlapping with each TF CUT&RUN peaks. Bottom, Heat plots showing SATB2 and other cortical TFs (LHX2, CUX1, BCL11A, NR2F1) binding peaks overlapping with each other and with CTCF ChIP-seq peaks. **d**, Western blots of co-immunoprecipitations from P0 cortices of *Satb2^+/+^* brains. Names on left indicate antibodies used for IP while names on right indicate antibodies used in the western blot. In: input, E: elution, IgG: IgG control elution.

To further investigate how SATB2 regulates genes associated with intelligence, we lifted over our mouse SATB2 ChIP-seq data to the human genome (hg38) and performed a comprehensive analysis to assess both the cross-species consistency and the functional conservation of SATB2 binding sites. First, to assess the concordance between human and mouse binding profiles, we obtained human SATB2 ChIP-seq data on NeuN^+^ cells isolated from dorsolateral prefrontal cortex from Loupe et al.^25^, and compared it against our mouse SATB2 ChIP-seq data mapped to the human genome (**Extended Data Fig 2b**). We observed high consistency between the datasets, as evidenced by the correlation of signal intensities in both the human SATB2 peaks and a background control set comprising an equivalent number of randomly sampled ENCODE candidate *cis*-regulatory elements (cCREs) (Spearman correlation: 0.48) (**Extended Data Fig 2b**). Second, to assess the functional importance of these binding sites, we analyzed these regions’ evolutionary conservation using 100-way vertebrate PhastCons scores (hg38). We compared the mean conservation scores of our identified SATB2 peaks against the same background control set. SATB2 peaks exhibit significantly higher evolutionary conservation compared to the background control (Wilcoxon two-sided rank-sum test*, p* < 2.2e-16) (**Extended Data Fig 2c**). This elevated conservation indicates that these genomic regions are under strong purifying selection.

We linked the SATB2 binding peak liftover to gene promoters using a published Proximity Ligation-Assisted H3K4me3 ChIP-seq (PLAC-seq) data set for human fetal cortical excitatory neurons^26,27^ (**Fig. 1b**). We observed that SATB2 potentially regulates the expression of many genes associated with intellectual deficiency including *AUTS2, CUX2, MEIS2,* and *TBR1* by directly binding to their promoters and/or enhancers (**Fig. 1b**). Collectively, these data reinforce the functional relevance of SATB2 in transcriptional regulation and support the translational validity of our mouse *Satb2* mutant model.

### Histone modifications are sensitive to SATB2 dosage

To determine how SATB2 dosage affects chromatin state, we performed CUT&RUN using cortices from P0 *Satb2^+/+^*, *Satb2^+/lacZ^*, and *Satb2^lacZ/lacZ^* mice and antibodies for SATB2 and active (H3K27Ac, H3K4me3) and repressive (H3K27me3) histone marks (**Extended Data Fig 3**). SATB2 CUT&RUN signal was robust in *Satb2^+/+^* cortices and was largely indistinguishable between *Satb2^+/+^* and *Satb2^+/lacZ^*samples, whereas SATB2 binding was absent in *Satb2^lacZ/lacZ^* cortices, consistent with loss of the SATB2 protein (**Extended Data Fig 3a**). In contrast, profiling of histone modifications revealed clear dosage-dependent shifts in chromatin state (**Extended Data Fig 3b**). When the histone peaks were stratified into regions with higher signal in *Satb2^+/+^* versus higher signal in *Satb2^lacZ/lacZ^*, wildtype (*WT*) enriched peaks showed a progressive reduction from *Satb2^+/+^*to *Satb2^+/lacZ^* to *Satb2^lacZ/lacZ^* whereas knockout (*KO*) enriched peaks increased in the opposite direction, with *Satb2^+/lacZ^*consistently exhibiting an intermediate pattern between *Satb2^+/+^* and *Satb2^lacZ/lacZ^* (**Extended Data Fig 3b**). However, the magnitude of this intermediate chromatin state was variable. For H3K4me3, *KO* enriched peaks were comparatively weak and showed only a modest increase across genotypes while *WT* enriched peaks only showed a slight decrease. For H3K27ac, the *WT* to heterozygous (*Het*) to *KO* transition was more pronounced, with a clear redistribution from *WT* enriched to *KO* enriched peaks in *Satb2^lacZ/lacZ^* cortices and an intermediate shift already evident in the *Satb2^+/lacZ^*. The strongest effect was observed for H3K27me3 where *Satb2^+/lacZ^* cortices displayed a balanced intermediate state across *WT* and *KO* enriched peak sets whereas *Satb2^lacZ/lacZ^* cortices showed a near complete inversion relative to *Satb2^+/+^* controls (**Extended Data Fig 3b**). These results indicate that SATB2 dosage has limited impact on genome-wide SATB2 occupancy at P0 but strongly influences chromatin state with *SATB2* heterozygous cortices adopting an intermediate epigenetic profile between *WT* and *KO*.

### Satb2 cooperates with other cortical transcription factors and BAF complex

We performed CUT&RUN and protein co-immunoprecipitation (co-IP) for SATB2, LHX2^28^, CUX1^29^, BCL11A^30,31^, and NR2F1^32,33^. These proteins are selected because they are known to regulate the development of intra-telencephalic (IT) projection neurons and have defined laminar specificity across L2/3-L6. The combination of snMultiome, CUT&RUN and co-IP analysis shows that they regulate common genomic targets with SATB2. Firstly, the E18 mouse brain snMultiome sequencing data show that the genes encoding these proteins are highly expressed in the SATB2-expressing cells (**Extended Data Fig. 4a**). Secondly, CUT&RUN peaks for SATB2, LHX2, CUX1, BCL11A and NR2F1 were enriched in H3K4me3, H3K27ac, and H3K27me3 modifications (**Extended Data Fig. 4b**), indicating that they all bind to gene promoters and potential enhancers. Based on the activity pattern of chromatin states and CTCF binding, the SATB2 peaks were categorized into three distinct clusters using *k*-means clustering (**Extended Data Fig. 4c**). Cluster 1 represents a small set of sites marked by high H3K4me3 and H3K27ac signals, consistent with active promoters. Cluster 2 is characterized by strong CTCF co-occupancy with moderate H3K27ac enrichment, and Cluster 3 displays moderate H3K27ac but low H3K4me3, a chromatin signature consistent with putative enhancers. Finally, CUT&RUN profiling and protein co-IP revealed that SATB2 co-occupies a substantial fraction of genomic sites with LHX2, CUX1, BCL11A, NR2F1, and CTCF, and physically associates with these factors *in vivo* (**Figs. 1c and 1d**). LDSC analysis showed that SATB2 binding regions exhibited the leading heritability enrichment for cognitive and neuropsychiatric traits among the transcription factors profiled, implicating SATB2 as a major regulator in the genetic architecture of human cognition and brain disorders (**Extended Data Fig. 4d**).

SATB2 was previously reported to inhibit transcription of the L5 PT genes, including *Bcl11b*, by recruiting the NuRD complex, which is mostly involved in transcriptional repression^10,11^. To investigate how SATB2 mediates transcriptional activation, we performed co-IP using antibodies for SATB2 and the BAF complex core component BRG1/SMARCA4 using *Satb2^+/+^* and *Satb2^lacZ/lacZ^* neocortices. Co-IP using a SATB2 antibody pulled down BRG1 and co-IP with a BRG1/BRM antibody (J1)^34^ pulled down SATB2 in the *Satb2^+/+^* cortical protein extract (**Extended Data Fig. 5a**). Immunostaining revealed co-expression of SATB2 and BRG1 in cortical projection neurons (**Extended Data Fig. 5b**). In a separate experiment^34^, a proteomic analysis of BAF interacting proteins in cultured cortical neurons identified SATB2 as one of the top hits co-purified with BAF at both basal and depolarized conditions (**Extended Data Fig. 5c**). These results together indicate that SATB2 cooperates with BAF chromatin remodeling complexes and activates transcription in cortical neurons.

### *Satb2* deficiency leads to differential expression of genes associated with ID

To determine how broadly cortical projection neurons are impacted by *Satb2* deletion and heterozygosity, we performed single nucleus RNA-sequencing (snRNA-seq) using dissected primary somatosensory cortex (SSp) from P28 *Satb2^+/fl^* (*control*), *Satb2^fl/lacZ^*(*het*), and *Satb2^fl/fl^*; *Emx1-Cre* (*cKO*) mice (*n* = 2 mice/genotype) (**Fig. 2 and Extended Data Fig. 6**). We focused on the SSp due to the extensive characterization of ExNs in this region in previous scRNA-seq analyses^9,35^, and because this is the dominant sensory processing area in rodent neocortices. After completing quality controls, 6,542 *Control*, 7,712 *Het,* and 8,920 *Satb2 cKO* cells were kept for analysis (**Extended Data Fig. 6a**). We annotated the cell clusters and identified cortical ExN, interneuron, oligodendrocyte precursor, oligodendrocyte, astrocyte, microglia, macrophage, endothelial, fibroblast, and striatal projection neuron clusters based on feature gene expression (**Extended Data Fig. 6b**). Since SATB2 is specifically expressed in ExNs, but not in progenitors or cortical interneurons, we focused our analysis on the ExN clusters (**Fig. 2a and Extended Data Fig. 6c**). In both *control* and *Het* cortices, we identified all IT neurons (L2/3 IT, L4 IT, L4/5 IT, L5 IT, and L6 IT), L5 near projecting (NP), L5 PT, L6 CT, and L6b cells (**Fig. 2a**). Spatial transcriptomic profiling with Slide-seq V2^36^ further confirmed the presence of all ExN clusters in *Satb2 Control* and *Het* cortices (**Extended Data Fig. 6d**).

**Figure 2:**
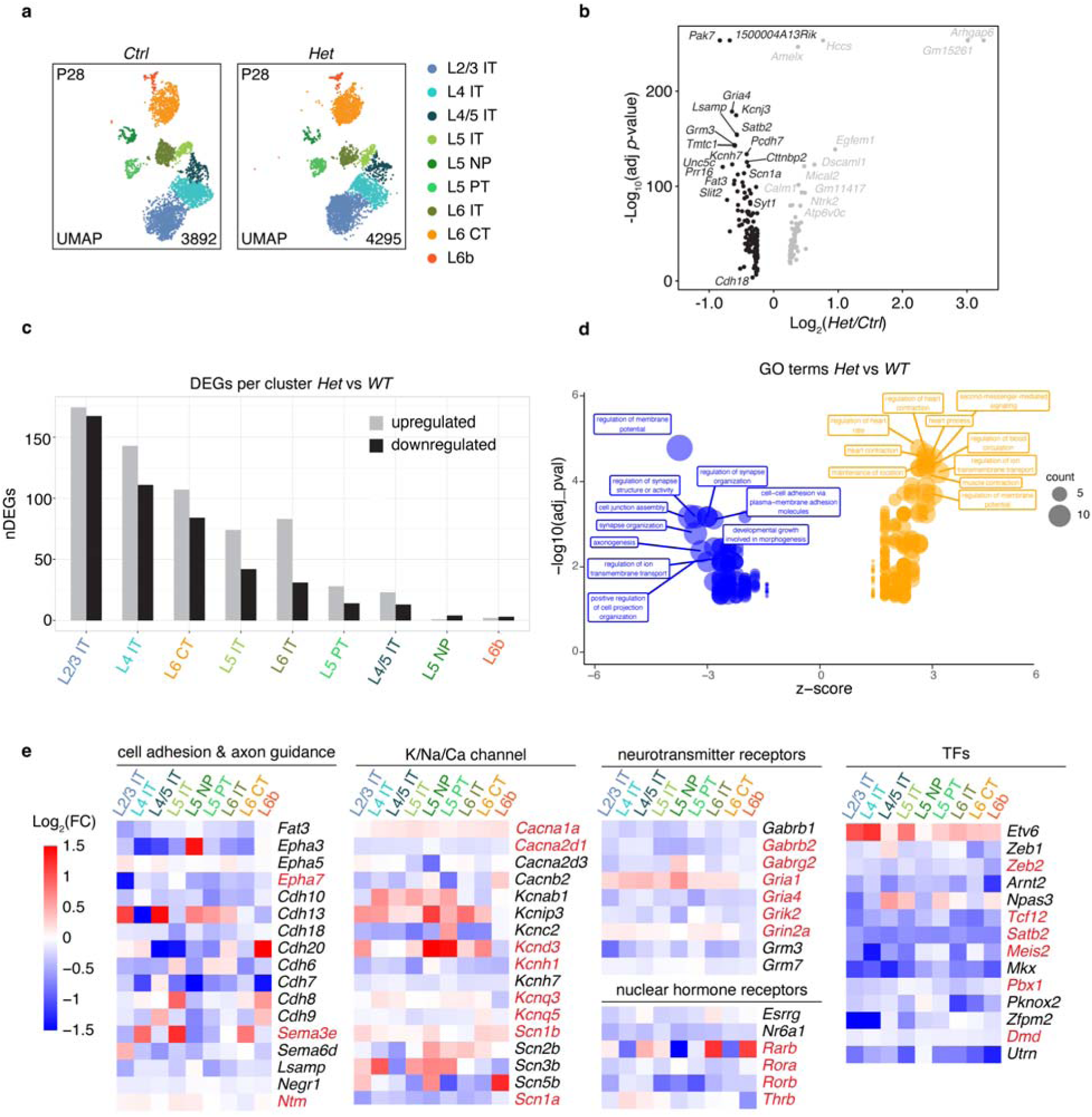
*Satb2* haploinsufficiency leads to mis-regulated gene expression, especially for genes associated with intelligence. **a**, UMAPs of ExN clusters in the SSp of *Satb2^+/fl^* (*Ctrl*) and *Satb2^fl/lacZ^* (*Het*) mice, colored according to cell types. **b**, Volcano plot showing upregulated and downregulated genes in combined ExN clusters in *Het* versus *Ctrl*. **c**, Numbers of DEGs between corresponding ExN clusters in *Satb2 Het* vs *Ctrl* samples. **d**, Gene ontology analysis of all DEGs in *Het* cortices versus *Ctrl* cortices, showing their z-score and unadjusted *p* values. Size of bubbles represents the number of DEGs associated with that term and the z-score represents the ratio of upregulated versus downregulated genes in the *Het* compared to *Ctrl*. Text boxes indicate the top 10 GO terms for up– or downregulated genes. **e**, Heatmaps of Log_2_(FC) of *Het* versus *Ctrl* separated by category of the protein-type each gene encoding for: cell and axon guidance, K/Na/Ca channel, neurotransmitter receptors, nuclear hormone receptors, and TFs. FC, Fold Change. ID genes are labeled in red.

We compared gene expression between corresponding *Satb2 control* and *Het* ExN clusters and identified hundreds of DEGs in the *Satb2 Het* neurons, many of which are cell-type-specific (**Fig. 2b-c and Extended Data Fig. 6e)**. Notably, the expression levels of the genes that were upregulated and downregulated genes in *Satb2 Het* were further altered in *Satb2 cKO* (**Extended Data Fig. 6f**), consistent with the increased phenotype severity in *Satb2 cKO*. The *Het* DEGs were associated with GO terms including regulation of membrane potential, synapse organization, retinal ganglion cell axon guidance, regulation of synapse structure or activity, cell junction assembly, etc (**Fig. 2d**). The most significant DEGs in the *Satb2 Het* ExNs include cell adhesion and axon guidance molecules, ion channels, neurotransmitter receptors, and transcription factors (**Fig. 2e**).

In the *Satb2 cKO* cortices, L5 PT and all IT clusters were lost, and 3 novel clusters were present (**Extended Data Figs. 6 and 7**). We determined the relationship between the novel ExN clusters and the missing neuronal cell types in the *Satb2 cKO* cells. To achieve this, we mapped gene expression features of each ExN cluster in control mice onto the *Satb2 cKO* ExN clusters, as well as the gene expression features of the 3 novel neuronal clusters in *Satb2 cKO* mice back onto the control ExN clusters (**Extended Data Figs. 7a-b**). This revealed that the *Satb2* cKO-1 and cKO-3 clusters resembled mutant L2/3 ITs, and the cKO-2 cluster resembled mostly deep-layer ITs and L5 PTs (**Extended Data Fig. 7b**). Further analysis of DEGs along with spatial transcriptomic profiling with Slide-seq V2 shows severe mis-regulation of canonical cortical development genes resulting from the lack of SATB2 (**Extended Data Fig. 7c-d**). Although L5 NP, L6 CT and L6b neurons were still present, differential gene expression analysis revealed hundreds of DEGs in each subtype when compared to the respective clusters in control mice (**Extended Data Fig. 7e**). Similar to the *Het, cKO* DEGs are associated with GO terms including synapse organization, axonogenesis, regulation of membrane potential, and cognition, among others (**Extended Data Fig. 7f**).

To examine changes in ID and intelligence related genes in *Satb2 Het*, we performed single-cell disease relevance score (scDRS) analysis^37^ to associate ID and intelligence related genes with cortical cell types using two risk gene sets from different GWAS studies (https://panelapp.genomicsengland.co.uk/panels/285/ and Savage et al., 2018^37^) (**Fig. 3**). The scDRS analysis significantly associated ID related genes with cortical ExN, interneuron, and astrocyte clusters and intelligence related genes with ExN clusters in the control cells, indicating that the expression of these genes is normally enriched in these clusters. However, this cell-type enrichment was no longer significant for most clusters in the *Satb2 Het* samples (**Fig. 3**). Consistent with the scDRS analysis result, many of these DEGs identified in the ExNs in the *Satb2 Het* mice were ID genes identified in GWAS studies, such as *Kcnd3*, *Gabrb2*, *Meis2*, *Sema3e*, *Gria1* and *Rorb* (**Fig. 2e**). These results indicate that heterozygous *Satb2* loss-of-function mutation leads to wide-spread mis-regulation of genes associated with intelligence and ID, and the disruption, but not loss, of ExN identities.

**Figure 3:**
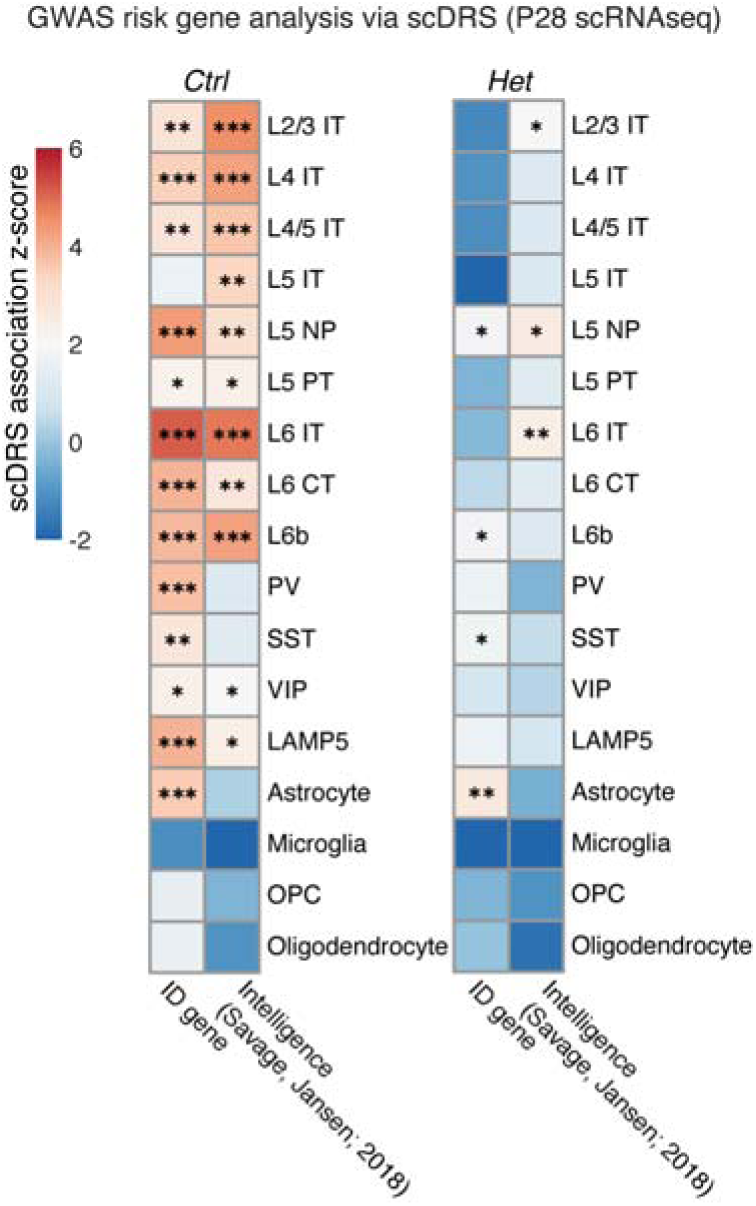
*Satb2* haploinsufficiency increases the risk of dysregulation of genes associated with ID. Identification of intellectual disability (ID)– and intelligence-relevant cell types based on the *Ctrl* and *Het* snRNA-seq data using scDRS analysis. (see Methods). \**p* < 0.05; \*\**p* < 0.01; \*\*\**p* < 0.001. Exact *p* values are provided in **Supplementary Table 2**.

### Defective cortical architecture in *Satb2^+/lacZ^* mice

To investigate how the mis-regulated gene expression in *Satb2^+/lacZ^*mice affects cortical development, we first performed western blots on *Satb2^+/+^*, *Satb2^+/lacZ^*, and *Satb2^lacZ/lacZ^* P0 cortices to assess the relative amount of SATB2 protein in each cortex (**Extended Data Fig. 8a-b**). *Satb2^+/lacZ^* cortices expressed about half the amount of SATB2 protein compared to the wildtype cortices, and no SATB2 protein was detected in the *Satb2^lacZ/lacZ^* cortex. We also show that early embryonic SATB2 expression is restricted to cortical ExNs and is not present in progenitor cells by immunostaining for SATB2, TBR1, and TBR2 from E11.5 to P0 (**Extended Data Fig. 8c**). We compared *Satb2^+/+^* and *Satb2^+/lacZ^* brains collected at P1, P4, P7, and P28 (**Fig. 4a-d**). Immunostaining in the SSp for markers of different cortical ExN subtypes and nuclear staining did not show a significant change in the *Satb2^+/lacZ^* cortices at P1. From P4 to adulthood, the thickness of L1 was increased, and the thicknesses of L4 and L6 were reduced, while the L5 thickness was not significantly affected in the *Satb2^+/lacZ^* cortices (**Fig. 4a-b**). TBR1 was expressed in L6 CTs, L6 ITs, L5/6 NPs, and L2/3 ITs, while TLE4 was specifically expressed in L6 CTs and L5/6 NPs^8^. In the *Satb2^+/lacZ^* cortices, the number of TBR1^+^TLE4^-^ L6 ITs was significantly reduced from P4 to adulthood across the medial-lateral and anterior-posterior axes of the cerebral cortex, while no significant changes in the numbers of the TLE4^+^ and TBR1^+^TLE4^+^ L6 CT and L5/6 NP neurons were observed (**Figs. 4c-d**). Immunostaining for BCL11B at P1, P4, and P7 revealed no significant changes in the numbers of BCL11B^+^ L5 PTs in *Satb2^+/lacZ^* cortices (**Fig. 4e-f**).

**Figure 4:**
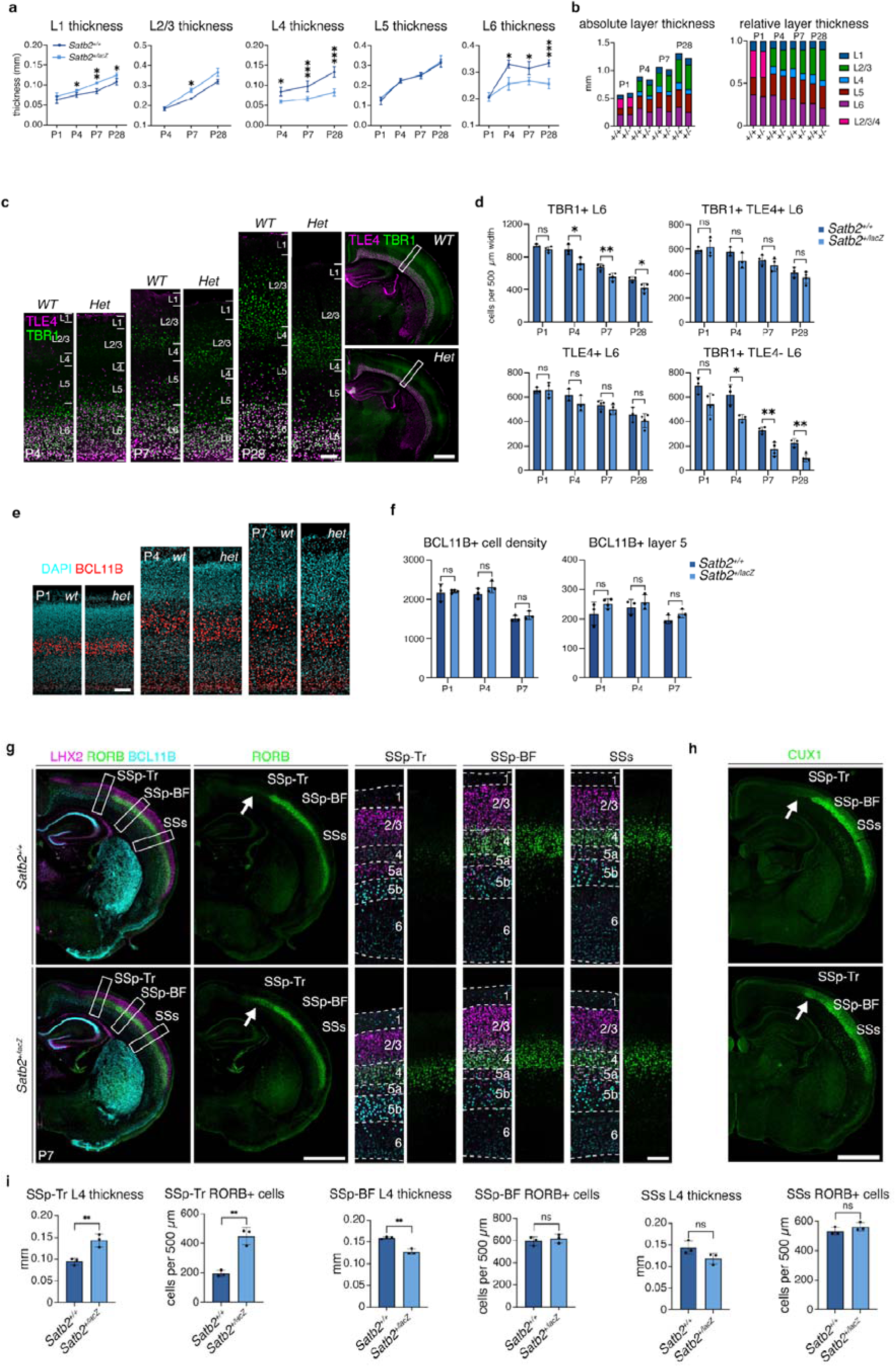
*Satb2^+/lacZ^*mice show altered cortical lamination. **a**, Quantification of cortical layer thickness over postnatal ages P1, P4, P7 and P28 in *Satb2^+/+^* and *Satb2^+/lacZ^* brains. Because L4 and L2/3 neurons are still migrating at P1, L4 and L2/3 measurements are not shown at P1. (P1: *n* = 3 *Satb2^+/+^*, *n* = 4 *Satb2^+/lacZ^* mice; P4: *n* = 3 mice per genotype; P7: *n* = 3 mice per genotype; P28: *n* = 4 mice per genotype. Detailed statistics in **Supplementary Table 1**). **b**, Absolute and relative layer thickness quantifications colored by layer. **c**, Immunostaining of TBR1 and TLE4, and DAPI labeling in P4, P7, and P28 brains. Scale bars: low mag, 1 mm; high mag, 100 µm. **d**, Quantification of TBR1^+^, TLE4^+^, TBR1^+^TLE4^+^ and TBR1^+^TLE4^-^ cells in L6 of 500-µm wide matching SSp regions. (P1: *n* = 3 *Satb2^+/+^*, *n* = 4 *Satb2^+/lacZ^*; P4: *n* = 3 per genotype; P7: *n* = 4 per genotype; P28: *n* = 3 *Satb2^+/+^*, *n* = 4 *Satb2^+/lacZ^*. Detailed statistics in **Supplementary Table 1**) **e**, Immunostaining of BCL11B and DAPI in P1, P4, and P7 cortices, Scale bar: 100 µm. **f**, Quantification of the L5 BCL11B^+^ cell number and cell density (cells per mm^2^) in 500-µm wide sections at P7 in matching SSp regions. (P1: *n* = 3 *Satb2^+/+^*, *n* = 4 *Satb2^+/lacZ^* mice; P4: *n* = 3 mice per genotype; P7: *n* = 3 mice per genotype. Detailed statistics in **Supplementary Table 1**). **g**, Immunostaining for LHX2, RORB, and BCL11B in P7 *Satb2^+/+^*and *Satb2^+/lacZ^* brains. White arrows highlight the SSp-Tr regions. Scale bars: low mag, 1 mm; high mag, 100 µm. **h**, Immunostaining for CUX1 in P7 *Satb2^+/+^* and *Satb2^+/lacZ^* brains. White arrow highlights the SSp-Tr regions. Scale bar 100 µm. **i**, Quantifications of L4 thickness and RORB^+^ cells per 500-µm wide brain section of indicated region. (*n* = 3 brains per genotype. Detailed statistics in **Supplementary Table 1**). For all graphs: error bars represent ± standard deviation (SD), ns, not significant; \**p* < 0.05, \*\**p* < 0.01, \*\*\**p* < 0.001.

We examined L2-4 ITs using antibodies for RORB, CUX1 and LHX2 in P7 *Satb2^+/+^* and *Satb2^+/lacZ^*cortices (**Fig. 4g-i**). In the *Satb2^+/lacZ^* brains, RORB^+^ cells in L4 and the thickness of L4 were significantly increased in the SSp trunk (SSp-Tr) region, while in primary somatosensory cortex barrel field (SSp-BF) there was a reduction in L4 thickness (**Fig. 4g,i**). In the secondary somatosensory region (SSs), no change was observed in L4 thickness or RORB^+^ cells. Like RORB, CUX1 expression was increased in the SSp-Tr (**Fig. 4h**). The changes in RORB and CUX1 expressions in L2-4 suggest defective somatosensory patterning in *Satb2^+/lacZ^* cortices.

We also examined cortical development defects in the prefrontal and motor cortical regions. In the medial prefrontal cortex (mPFC), immunostaining for TBR1 and TLE4 revealed no significant differences in the numbers of TBR1^+^ or TBR1^+^TLE4^-^ L6 IT neurons, TLE4^+^ corticothalamic neurons, or TBR1^+^TLE4^+^ L6 neurons between *Satb2^+/+^* and *Satb2^+/lacZ^* P7 cortices (**Extended Data Fig. 9a**). These findings indicate that reduced SATB2 dosage does not disrupt L6 ExN composition in the mPFC. In contrast, analysis of the primary motor cortex (MOp) revealed pronounced and broad deficits in L6 projection neurons in *Satb2^+/lacZ^* mice. Quantification showed significant reductions in the numbers of TBR1^+^TLE4^-^ L6 IT neurons, TLE4^+^ CT neurons, and TBR1^+^TLE4^+^ L6 neurons compared to wildtype controls (**Extended Data Fig 9b**). This aligns with findings that SATB2 is a critical regulator of primary sensorimotor area development^38^.

To examine if the laminar organization defects are unique to the *Satb2^lacZ^*allele, or if they occur with other *Satb2* LOF alleles, we examined cortical ExNs in the *Satb2^+/fl^*; *Emx1-Cre* mice and observed similar defects (**Extended Data Fig. 10**). Thus, heterozygosity of *Satb2* loss-of-function mutation leads to defective cortical laminar organization.

### *Satb2* haploinsufficiency results in defective IT axons

Because SATB2 regulates gene expression across the major ExN cell-types, we examined ExN projection changes due to *Satb2* haploinsufficiency. Immunostaining with an L1 antibody revealed that in *Satb2^+/lacZ^* mice, the corpus callosum length was significantly reduced along the anterior-posterior axis at P7, but L1 and PKCG staining did not reveal obvious changes in corticothalamic or corticospinal axons (**Figs. 5a-b**).

**Figure 5:**
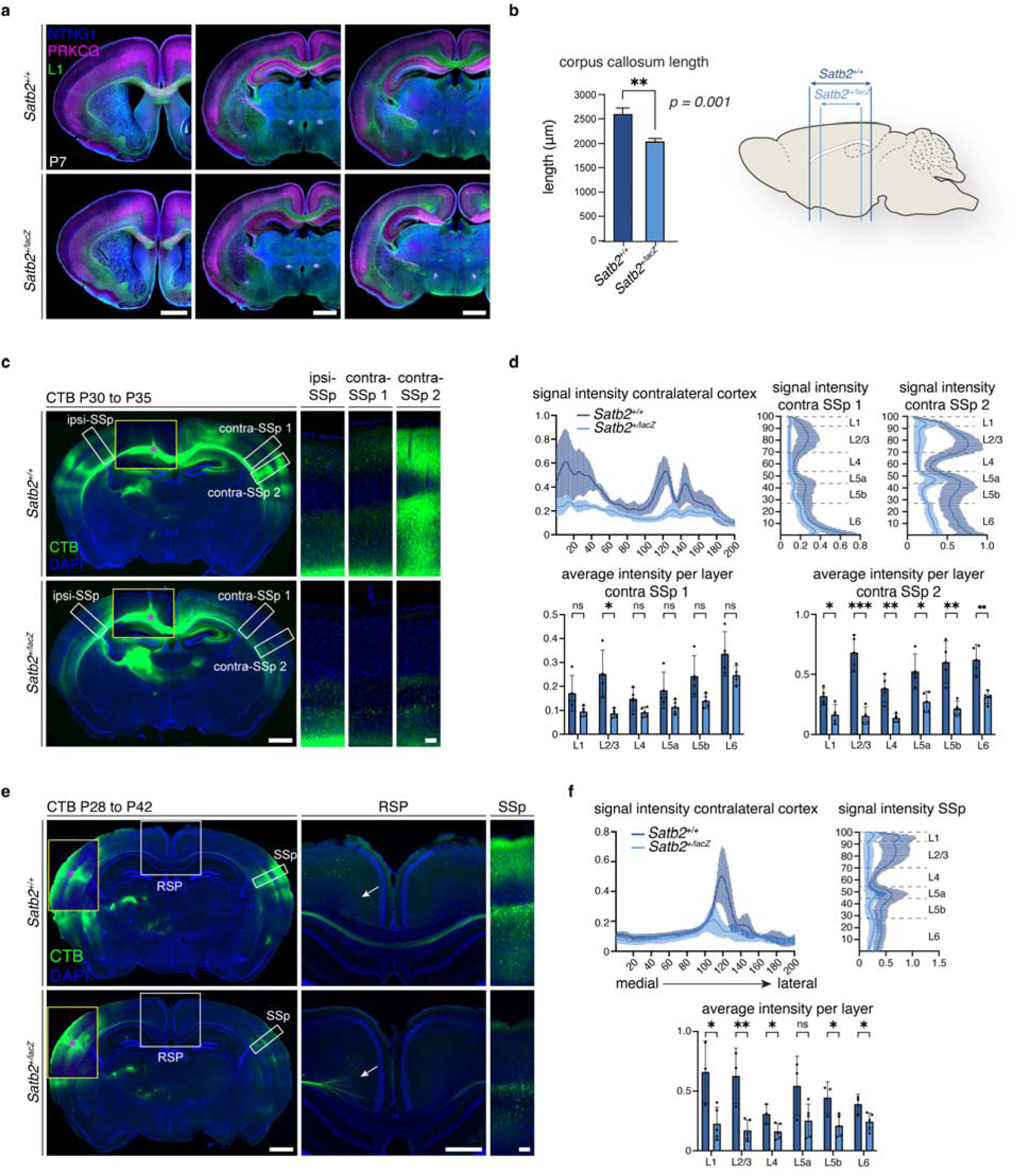
Intra-telencephalic projection neurons and axons are altered in *Satb2^+/lacZ^* brains. **a**, Immunostaining of L1, NTNG1, and PRKCG in P7 *Satb2^+/+^* and *Satb2^+/lacZ^* brains. Each row shows progressively more posterior sections. Scale bars, 1 mm. **b**, Quantification of corpus callosum length along the anterior-posterior axis in fixed brains at P7 (*n* = 10 brains per genotype, *p = 0.001*, unpaired *t*-test). Diagram indicates anterior and posterior boundaries for each genotype. **c**, CTB tracing in brains with CTB injected in the CC at P30 and analyzed at P35, counterstained with DAPI. Yellow boxes indicate lower-exposure views of the injection sites; white boxes show magnified areas in ipsilateral (ipsi-SSp) and contralateral (contra-SSp 1, 2) primary somatosensory cortex. Asterisk marks injection site. Scale bars: low magnification, 1 mm; high magnification, 100 µm. **d**, Quantifications of CTB fluorescence intensity from **c**, normalized to injection site intensity. Top left: signal along the medial-lateral axis of the contralateral cortex. Top right, normalized CTB signal along the apical-basal axis in contralateral SSp (regions 1 and 2). Bar graphs show mean layer-specific fluorescence intensity in contralateral SSp regions 1 and 2. (*n* = 4 *Satb2^+/+^*, 5 *Satb2^+/lacZ^* brains. See **Supplementary Table 1** for statistics). **e**, CTB tracing following SSp injection at P28 and analysis at P42. DAPI staining in blue. Yellow boxes indicate injection sites, white boxes show magnified retrosplenial area (RSP) and contralateral SSp. White arrows highlight defasciculating axons observed in *Satb2^+/lacZ^* but not *Satb2^+/+^* cortices. Asterisks mark injection sites. Scale bars: low magnification full section, 1 mm; high magnification RSP, 500 µm; high magnification contralateral SSp, 100 µm. **f**, Quantifications of CTB fluorescence from **e**, normalized to injection site intensity. Top left: medial-lateral distribution in contralateral cortex. Top right, apical-basal distribution across six cortical layers of contralateral SSp. Bar graphs show mean layer-specific fluorescence (*n* = 4 brains per genotype, See **Supplementary Table 1** for statistics).

We then systematically examined callosal, subcerebral, and corticothalamic axons using anterograde and retrograde tracing (**Figs. 5c-f, and Extended Data Fig. 11**). We injected cholera toxin subunit B (CTB) directly into the corpus callosum of one cortical hemisphere near the midline, which anterogradely labeled passing axons to their targets and retrogradely labeled the cell bodies of the neurons projecting axons through the injection site (**Figs. 5c-d**). We observed CTB-labeled cells and axons in L2/3 and in L5/6 in the ipsilateral and contralateral cortices in *Satb2^+/+^* mice along the medial-lateral cortical axis. In *Satb2^+/lacZ^* cortices, there was a near complete absence of signal in L2/3 with reduced labeling of cells and axons in L5/6 in both ipsilateral and contralateral cortex (**Figs. 5c-d**), indicating severe defects in both ipsilateral and callosal axons. To further investigate callosal projections, we injected CTB into the SSp of one cortical hemisphere (**Fig. 5e**). In the *Satb2^+/+^* mice, labeled axons formed a tight bundle in the corpus callosum, crossing the midline and innervating the contralateral SSp region across all layers except L4. In the *Satb2^+/lacZ^* brains, labeled callosal axons defasciculated as they approached the midline. Significantly less CTB signal was measured in the deep layers of the contralateral SSp, and most notably, there was a severe reduction of signal measured in L2/3 (**Figs. 5e-f**). Injecting CTB into the ventroposterior medial thalamic nucleus to retrogradely label L6 CTs (**Extended Data Fig. 11a**), or into the cerebral peduncle to label L5 PTs (**Extended Data Fig. 11b**) indicate that L6 CT or L5 PT projections were not noticeably affected in the *Satb2^+/lacZ^* cortices.

To investigate layer-specific intra-telencephalic axon projections, we performed cell type-specific axon tracing by injecting Cre-dependent AAV9-FLEX-GFP virus into the SSp of *Satb2^+/+^* and *Satb2^+/lacZ^* mice that carried a *Cux2^CreER^* (**Extended Data Fig. 11c**), *Ntsr1-Cre* (**Extended Data Fig. 11d**), or *Tlx3-Cre* (**Extended Data Fig. 11e**) allele. The results showed that in *Satb2^+/lacZ^* mice, callosal axons from *Cux2^CreER^*^+^ L2/3 IT neurons defasciculated near the midline and had dramatically reduced innervation of the contralateral SSp (**Extended Data Fig. 11c**). Axons from *Tlx3-Cre^+^*L5 IT neurons extended into contralateral SSp (**Extended Data Fig. 11e**) but showed reduced projection into the temporal association area (TeA). Most *Ntsr1-Cre+* neurons were L6 CTs and projected axons into the thalamus, but a minor population of them extended callosal axons, which were retrogradely labeled by the AAV9-FLEX-GFP virus in the SSp of *Satb2^+/+^* contralateral hemisphere (**Extended Data Fig. 11d**). In the *Satb2^+/lacZ^* mice, no retrograde labeled L6 ITs were observed in the contralateral SSp region (**Extended Data Fig. 11d**).

These findings reveal that *Satb2* haploinsufficiency leads to a severe reduction in intra-telencephalic projections, especially for L2/3 ITs and L6 ITs, while L6 corticothalamic and L5 subcerebral axons are spared.

### Reduced activity and defective local circuits for *Satb2^+/lacZ^* cortical projection neurons

To examine if *Satb2* haploinsufficiency affects neuronal physiology, we performed whole-cell patch-clamp recordings in acute brain slices on retrogradely labeled L2/3 IT and L5 IT neurons in SSp of P28 *Satb2^+/+^*and *Satb2^+/lacZ^* mice (**Fig. 6 and Extended Data Fig. 12**). Compared to neurons in *Satb2^+/+^* cortices, *Satb2^+/lacZ^*L5 IT neurons exhibited significantly increased rheobase current and action potential (AP) half width (**Fig. 6b**). Measuring AP firing in response to depolarizing current steps revealed that the same current steps led to overall reduced AP firing in IT neurons in *Satb2^+/lacZ^* mice, in addition to significantly decreased miniature excitatory postsynaptic current (mEPSC) amplitude and frequency (**Fig. 6c-e for L5 IT neurons and Extended Data Fig. 12a for L2/3 IT neurons**). These data suggest that L2/3 and L5 neurons in the SSp of *Satb2^+/lacZ^* mice are intrinsically less excitable.

**Figure 6:**
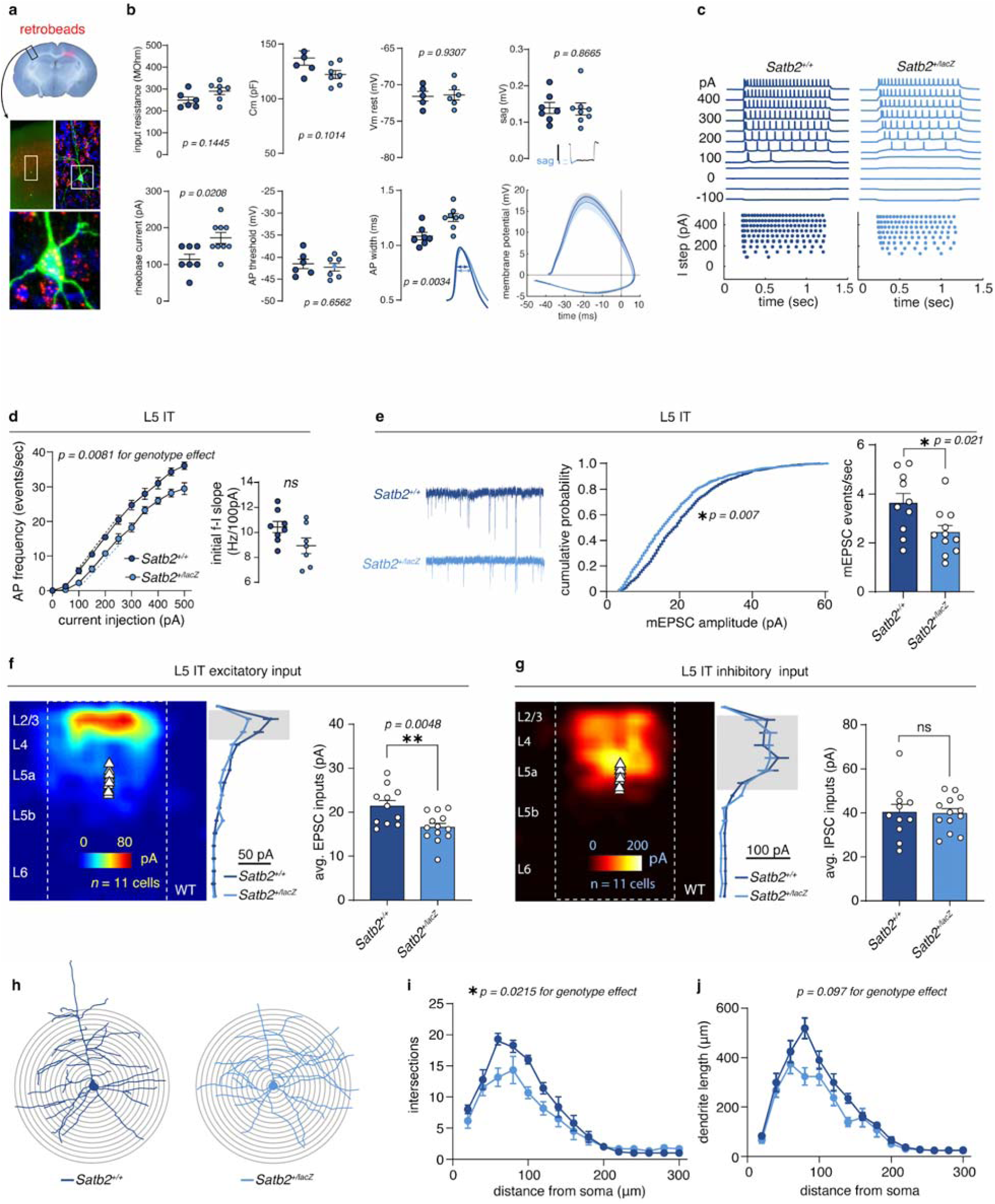
Electrophysiological changes of *Satb2^+/lacZ^* L5 IT neurons. **a**, Schematic of retrobead tracing used to identify IT neurons for electrophysiology recordings. **b**, Intrinsic membrane properties of L5 IT neurons measured by whole cell patch clamp, including input resistance (*n* = 6 *WT*, 7 *Het*), membrane capacitance (*n* = 6 *WT,* 7 *Het*), resting membrane potential (*n* = 5 *WT*, 6 *Het*), *s*ag (*n* = 7 *WT*, 8 *Het*), rheobase (*n* = 7 *WT*, 9 *Het*), action potential (AP) threshold (*n* = 6 *WT*, 7 *Het*), and AP width (*n* = 7 *WT*, 8 *Het*). All comparisons by two-tailed Mann-Whitney U test. **c**, Representative AP firing traces from L5 IT neurons during stepwise current injections (f-I protocol). **d**, Average firing frequency-current (f-I) curves and and initial f-I slopes for *Satb2^+/+^*and *Satb2^+/lacZ^* L5 IT neurons. (f-I curves: *n* = 4 *WT*, 4 *Het*, Two-Way Anova; f-I slopes: *n* = 8 *WT*, 7 *Het,* two-tailed Mann-Whitney U test). **e**, Representative mEPSCs, cumulative distributions, and event frequencies in *Satb2^+/+^* and *Satb2^+/lacZ^*L5 IT neurons (*n* = 10 *WT*, 11 *Het,* K-S test for cumulative distributions; two-tailed Mann-Whitney U test for frequency). **f-g**, LSPS maps of EPSC (**f**) and IPSC (**g**) inputs. Color maps show averaged responses from 5-6 *Satb2^+/+^* neurons; grouped synaptic inputs (mean ± SD) are summarized by cortical layer. Bar graphs compare total EPSC and IPSC inputs between *Satb2^+/+^* and *Satb2^+/lacZ^* neurons (*n* = 11 *WT,* 13 *Het*, two-tailed Mann-Whitney U test). **h**, Representative Sholl analyses of L5 IT neurons from *Satb2^+/+^* and *Satb2^+/lacZ^* cortices. **i**, Quantified Sholl intersection profiles of L5 IT *Satb2^+/+^* and *Satb2^+/lacZ^* neurons (*n* = 5 *WT*, 6 *Het).* Genotype effect: F_(1,9)_ = 7.706, *p = 0.0215*; Two-Way Anova). **j**, Dendrite length profiles of L5 IT neurons from *Satb2^+/+^*and *Satb2^+/lacZ^* cortices (*n* = 5 *WT*, 6 *Het* neurons). Genotype effect: F_(1,9)_ = 3.366, *p = 0.0997,* Two-Way Anova.

We applied laser-scanning photostimulation (LSPS), to provide a functional readout of local circuit connectivity with cellular resolution^39–41^, and glutamate uncaging^39,42^ to map circuit connectivity in the SSp of P28 control and *Satb2^+/lacZ^*mice (**Figs. 6f-g and Extended Data Fig. 12b**). We recorded intracortical excitatory connectivity for both L5 ITs and L2/3 ITs in the SSp in control and *Satb2^+/lacZ^* brain slices. Compared to the *Satb2^+/+^*cells, L5 ITs in the *Satb2^+/lacZ^* brain slices received significantly reduced excitatory input from L2/3 (**Fig. 6f**); L2/3 ITs showed no change in excitatory input from L5 (**Extended Data Fig. 12b**). Inhibitory inputs to L5 ITs and to L2/3 ITs were unchanged in the *Satb2^+/lacZ^* brain slices (**Fig. 6g and Extended Data Fig. 12b**), consistent with absence of SATB2 expression in cortical interneurons.

To determine whether these functional changes were accompanied by alterations in dendritic morphology, we performed Sholl analysis on the recorded L5 IT and L2/3 IT neurons (**Fig. 6h-j for L5 and Extended Data Fig. 12c-e for L2/3**). Dendrites of L5 IT neurons in *Satb2^+/lacZ^*mice showed a significant reduction in complexity with fewer dendritic intersections compared to *Satb2^+/+^* controls (**Fig. 6i**). Total dendritic length of L5 IT neurons was not significantly altered (**Fig. 6j)**. Similarly, dendrites of L2/3 IT neurons in *Satb2^+/lacZ^* mice showed a significant reduction in complexity with fewer dendritic intersections compared to *Satb2^+/+^*controls and no difference in length **(Extended Data Fig. 12c-e**). These data indicate that *Satb2* haploinsufficiency results in decreased L2/3 IT and L5 IT dendritic arborization along with reduced intrinsic excitability in the SSp.

### Abnormal somatosensory processing in *Satb2^+/lacZ^* mice

SAS patients display a broad spectrum of behavioral abnormalities^43^. Data from the SAS registry (accessed 2/2026) revealed various sensory-related abnormalities, including high tolerance to painful stimulation, skin or nail picking, mouthing and chewing of non-food items, and altered sensitivity to noise, touch, or certain textures. In addition, tactile disturbances are common in neurodevelopmental disorders^44^. Mice distinguish different textures using their whiskers, which relies on sensory processing in the SSp. In rodents, the texture information is encoded by L2/3 ExNs^45^ and is integrated into and processed within L5 ExNs whose activities eventually guide the behavioral output^46^. Using a texture preference test, we determined whether *Satb2* haploinsufficiency affects somatosensory information processing (**Figs. 7a-d and Extended Data Figs. 13a-d**). *Satb2^+/+^* and *Satb2^+/lacZ^* mice were presented with two textures, and the percentages of time they spent exploring each texture were measured. The *Satb2^+/+^* mice spent more time with the 120-grit sandpaper (120-grit) compared to the smooth (Sm) texture (**Fig. 7a**). Consistent with the defects in intrinsic neuronal physiology and local cortical circuitry in SSp, the *Satb2^+/lacZ^* mice did not differentiate between Sm and 120-grit textures (**Fig. 7a**). Bilateral whisker trimming eliminated the preference between Sm and 120-grit in *Satb2^+/+^* mice and had no effect in *Satb2^+/lacZ^* mice (**Fig. 7b**), confirming this behavior is dependent on sensory input from whiskers. This behavior phenotype was observed in both male and female *Satb2^+/lacZ^* mice **(Fig. 7c-d)**.

**Figure 7:**
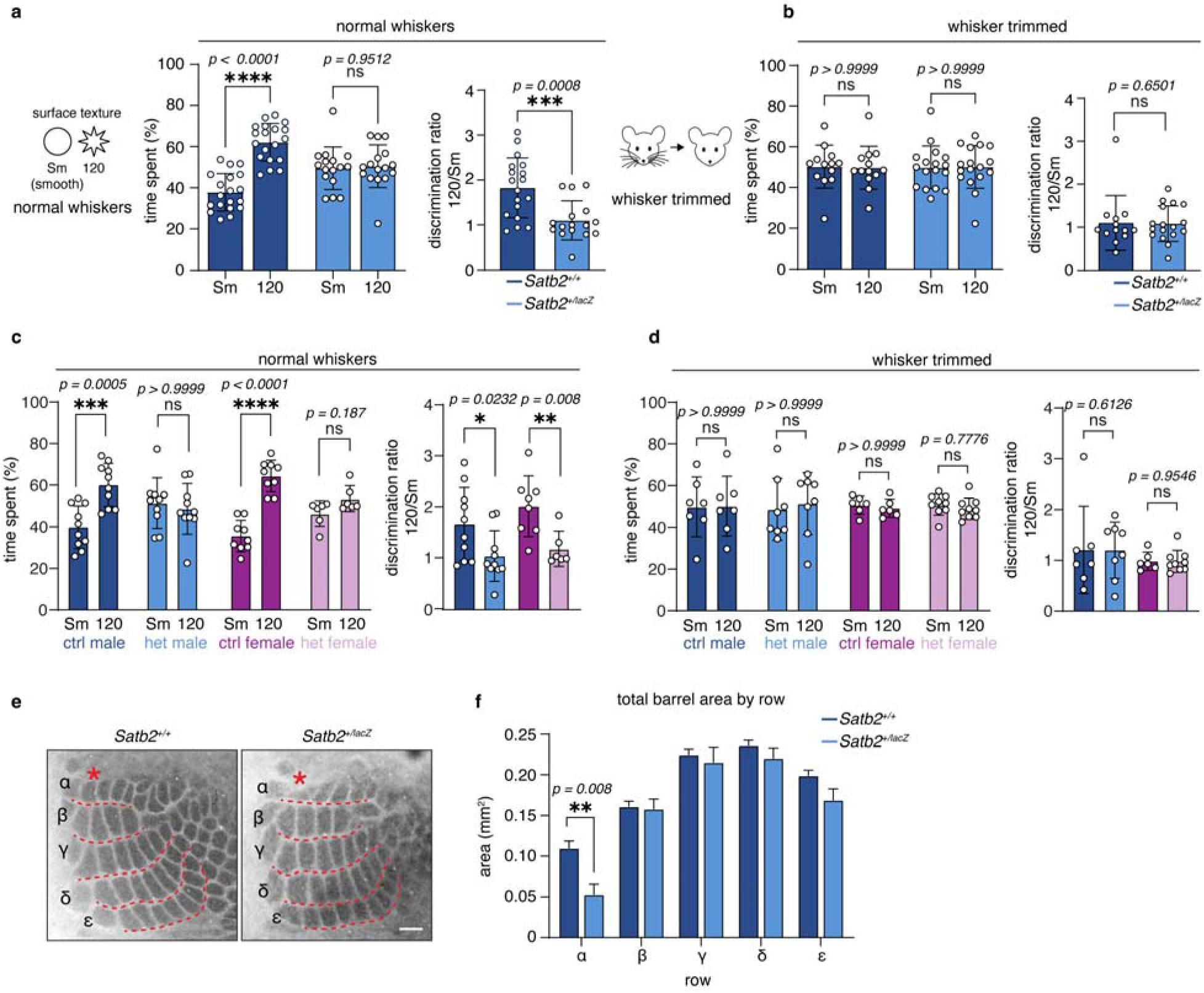
Decreased SATB2 dosage alters the function of primary somatosensory cortex. **a**, Texture discrimination between smooth (Sm) and 120-grit (120) sandpaper in *Satb2^+/+^* (ctrl) and *Satb2^+/lacZ^* (het) mice, quantified as percentage of time spent per texture and discrimination ratio (*n* = 19 ctrl,16 het) Time spent: Two-Way Anova with Bonferroni correction; discrimination ratio: Mann-Whitney test. **b**, Texture discrimination following whisker trimming, quantified as above (*n* = 13 ctrl, 17 het). Time spent: Two-Way Anova with Bonferroni correction; discrimination ratio: Mann-Whitney test. **c**, Texture discrimination with intact whiskers, analyzed by sex and genotype. Percentage of time spent per texture and the discrimination ratio are shown (n = 10 ctrl male, 9 ctrl female, 10 het male, and 6 het female). Time spent: Two-Way Anova with Bonferroni correction; discrimination ratio: Mann-Whitney test. Error bars represent ± SEM. **d**, Texture discrimination with whisker trimming, analyzed by sex and genotype (*n* = 10 ctrl male, 9 ctrl female, 10 het male, and 6 het female). Time spent: Two-Way Anova with Bonferroni correction; discrimination ratio: Mann-Whitney test. Error bars represent ± SEM. **e**, Cytochrome c oxidase staining of flattened P7 cortices from *Satb2^+/+^* and *Satb2^+/lacZ^*brains. Star illuminates missing barrels. Scale bar: 500 µm. **f**, Quantifications of total barrel area summed by barrel row in *Satb2^+/+^* and *Satb2^+/lacZ^*brains (*n* = 4 WT and Het). Two-Way Anova with Šídák’s multiple comparisons test.

To determine if decreased SATB2 dosage in defined L5 ExN subpopulations disrupts somatosensory discrimination, we performed texture preference assays using two L5 Cre-dependent SATB2 heterozygous models, *Satb2^+/fl^*; *Rbp4-Cre* and *Satb2^+/fl^*; *Tlx3-Cre*. *Satb2^+/fl^*; *Rbp4-Cre* mice, which targeted L5b ExNs, exhibited a loss of texture preference consistent with the phenotype observed in *Satb2^+/lacZ^* mice and failed to show a preference between Sm and 120-grit textures (**Extended Data Fig. 13a-b**). This impairment was consistent across both male and female mice. In contrast, *Satb2^+/fl^*; *Tlx3-Cre* mice, which targeted L5a population, show a stark reversal of texture preference, spending significantly more time interacting with the smooth texture relative to wildtype controls (**Extended Data Fig 13c-d**). This reversal in texture preference was seen in males but did not reach statistical significance in the female group. These results indicate that SATB2 heterozygosity in distinct L5 IT neuron subpopulations differentially affects texture-based sensory discrimination behavior.

We next examined barrel fields, a readily discernable SSp cortical structure composed of clustered thalamocortical afferents that relay somatosensory information from the whiskers, using cytochrome-c oxidase (CO) staining. Barrel organization was discernable in both control and *Satb2^+/lacZ^* brains. However, in *Satb2^+/lacZ^* cortices, the barrel field showed a disrupted organization, with barrels in the A1 row consistently missing (**Figs. 7e-f**).

### Abnormal pup calls in *Satb2^+/lacZ^* mice

One prominent feature of SAS is absent or severely reduced speech. To determine whether heterozygosity of *Satb2* loss-of-function mutation affects communication in mice, we recorded and analyzed isolation-induced ultrasonic vocalizations (USVs) in neonatal and early postnatal mice at P3, P7, and P11 (**Extended Data Fig. 13e-f**). A total of 72,253 calls were recorded from *Satb2^+/+^*, *Satb2^+/lacZ^*, and *Satb2^+/Fl^; Emx1^Cre/+^* mice. Calls were classified according to established criteria^47^, and multiple acoustic parameters were quantified.

*Satb2^+/lacZ^* pups showed significant alterations in call number, call length, median-, low-, and high-frequency of calls, slope, mean power, tonality, and peak frequency relative to *Satb2^+/+^* littermates, indicating impaired vocalization behavior (**Extended Data Fig. 13e-f**). Because SATB2 plays essential roles in both cortical ExN differentiation and craniofacial development, we next assessed whether these USV abnormalities could be attributed specifically to reduction of cortical SATB2. We recorded USVs from *Satb2^+/Fl^; Emx1^Cre/+^* pups, in which *Satb2* heterozygosity was restricted to cortical ExNs. In contrast to *Satb2^+/lacZ^* pups, *Satb2^+/Fl^; Emx1^Cre/+^* pups did not exhibit significant differences in USV acoustic parameters compared to *Satb2^+/+^* controls.

These findings indicate that the vocalization deficits observed in *Satb2^+/lacZ^* pups might also arise from a non-cortical etiology, most likely related to craniofacial abnormalities. These results suggest that speech deficits seen in SAS patients with LOF variants arise from either cortical or peripheral developmental defects.

## Discussion

SAS is a neurodevelopmental disorder defined by severe intellectual disability and speech deficits^1,2,18^. Although the external manifestations of intelligence differ between mice and human, the molecular programs and circuit architectures that support perception and cognition are broadly conserved, and prior cortical studies of SATB2 have relied almost exclusively on homozygous loss-of-function mutants that do not reproduce patient genotype. By characterizing a *Satb2* heterozygous mouse that models the gene dosage seen in SAS, we establish a mechanistic framework in which SATB2 functions as a dose-sensitive chromatin regulator of postnatal cortical maturation. The resulting phenotypes are distinct from those of the homozygous null and more directly explain the cognitive and sensory features of SAS.

Genome-wide binding analyses place SATB2 preferentially at distal regulatory elements adjacent to genes associated with intelligence and neurodevelopmental disorders. SATB2 co-immunoprecipitates with multiple chromatin regulatory complexes, including the BAF and NuRD remodelers and the polycomb repressive complex 2 (PRC2) (**Extended Data Figs. 5d-f**), as well as transcription factors enriched in cortical ExNs. This is consistent with a model in which SATB2 acts as a central regulatory scaffold that coordinates transcriptional control. Critically, reduced SATB2 dosage does not redistribute binding (as seen in the knockout) but instead alters chromatin state at its target loci, positioning the heterozygous as an intermediate chromatin configuration between wild-type and null rather than a partial loss of site occupancy.

At P28, snRNA-seq and spatial transcriptomics revealed that excitatory neuron subtypes are correctly specified in *Satb2*^+/lacZ^ cortex, yet each subtype exhibits extensive, cell-type-specific transcriptional dysregulation. DEGs cluster in functional categories that directly predict the circuit-level phenotypes, including regulation of membrane potential, synapse organization and activity, axon guidance, and cell-cell junctions. Altered transcription of voltage-gated ion channels and neurotransmitter receptor subunits maps onto reduced intrinsic excitability and diminished mEPSC amplitude and frequency, while dysregulation of EPH receptors, cadherins, and semaphorins^38,48–51^ maps onto IT axon targeting and disrupted barrel cortex organization. Single-cell scoring against gene sets associated with intelligence and intellectual disability was reduced across ExN subtypes and – notably – also in non-ExN populations including interneurons, pointing to non-cell-autonomous consequences of SATB2 haploinsufficiency that may contribute to SAS features including intellectual disability and behavior abnormalities.

Disease relevance scoring using gene sets associated with intellectual disability and intelligence indicates that cell-type specific enrichment of these programs is diminished in the *Satb2^+/lacZ^* cortex, supporting the conclusion that reducing SATB2 dosage disrupts conserved genetic programs implicated in cognitive and neurodevelopmental disorders. Although SATB2 is primarily expressed in ExNs, dysregulated disease-associated gene programs were also detected in additional cell types, including interneurons, suggesting non-cell-autonomous interactions may further shape SAS phenotypes.

Unlike the embryonic fate defects of homozygous *Satb2* loss-of-function, the structural and functional phenotypes of *Satb2*^+/lacZ^ mice emerge postnatally and converge on failures of circuit maturation rather than lineage specification. IT axon projections are selectively and subtype-specifically impaired: callosal and intracortical connectivity of L2/3 and L6 IT populations is markedly reduced, L5 IT callosal projections are mis-targeted, while L6 CT and L5 PT projections are spared. These projection defects are accompanied by region-specific laminar changes, with large reductions of TBR1^+^ L6 neurons in SSp and MOp, but relative preservation in mPFC, indicating that cortical areas differ in their sensitivity to SATB2 dosage. Both L5 and L2/3 ExNs additionally show reduced dendritic arborization. Together these findings show that *SATB2* haploinsufficiency disrupts multiple partially redundant developmental programs whose combined effects manifest as failures in IT targeting, dendritic maturation, and local circuit assembly.

To capture the functional output of these circuit defects, we exploited whisker-dependent texture discrimination^46,52^, an assay that engages precisely the sensory cortex circuits disrupted in *Satb2*^+/lacZ^ mice. Whereas wild-type mice show a robust preference for the 120-grit texture, *Satb2*^+/lacZ^ mice show no preference. Conditional SATB2 reduction reveals that this behavioral phenotype is cell-type specific: *Satb2^+/fl^* mice with reduced SATB2 in L5 PT neurons lose texture preference, whereas the same reduction in L5 IT neurons produces a distinct reversal of preference toward the smooth texture. These divergent outcomes reinforce that *SATB2* haploinsufficiency produces cell-type-specific effects on circuit function and provide a disease-relevant functional endpoint consistent with the sensory abnormalities reported by SAS caregivers.

*Satb2*^+/lacZ^ pups also exhibit significant alterations in isolation-induced ultrasonic vocalizations (USVs), a rodent correlate of the speech deficits that define SAS, but these alterations were largely absent in *Satb2^+/fl^*; *Emx1-Cre* pups, in which SATB2 reduction is restricted to cortical excitatory neurons. This dissociation implies that the USV phenotype of the global heterozygote arises largely from non-cortical contributions, plausibly craniofacial or peripheral oromotor mechanisms consistent with the dysmorphic facial features and feeding/oromotor difficulties documented in SAS^1,2^ and supports a model in which *Satb2* haploinsufficiency shapes behavior through multiple developmental pathways.

Taken together, our findings provide an integrated molecular-to-behavior framework for *Satb2* haploinsufficiency that directly explains SAS, linking dose-sensitive transcriptional and chromatin alterations to changes in neuronal excitability, connectivity, and sensory discrimination. Because these phenotypes emerge postnatally, they define an early postnatal window in which restoring SATB2-dependent programs could plausibly alter the trajectory of cortical dysfunction in SAS. More broadly, the results underscore that faithful modeling of patient gene dosage, rather than constitutive loss of function, is essential for defining mechanisms and identifying therapeutic entry points in neurodevelopmental disorders.

## Acknowledgments

This study was supported by grants to B Chen (NIH R01MH094589 and R01NS089777), G Fishell (NIH R01NS081297 and R37MH071679), S Qiu (NIH R01MH128192, R01EY035138), L Chen (NIH R01NS115660), J Wu (NIH R01NS137584), and D Feldheim (NIH R21 EY032230). TS Finn was supported by NIH T32HD108079, S Wu was supported by NIH F32MH125464, G Servito was supported by CIRM Training Program in Systems Biology of Stem Cells EDUC4-12759. We would like to acknowledge technical support and training from Benjamin Abrams in the UC Santa Cruz Life Sciences Microscopy Center. Use of the Zeiss 880 confocal microscope was made possible by the award of NIH S-10 Instrumentation grant NIH S10OD23528.

## Author contributions

BC, TSF, JT, MD, SJW, LC, and SQ designed the research, TSF, JT, MD, SJW, GS, XM, JZ, EP, CG, HL, AS, AAS, and YS performed the experiments. TSF, JT, MD, SW, SQ, EP, SK, AF, SM and BC analyzed the data. TSF, JT, MD, SJW, GF, and BC wrote the manuscript. GS, ES, CG, AS, AAS, JW, YZ, LC and SQ edited the manuscript.

## Materials & Correspondence

Correspondence and material requests should be addressed to.

**Extended Data Figure 1:**
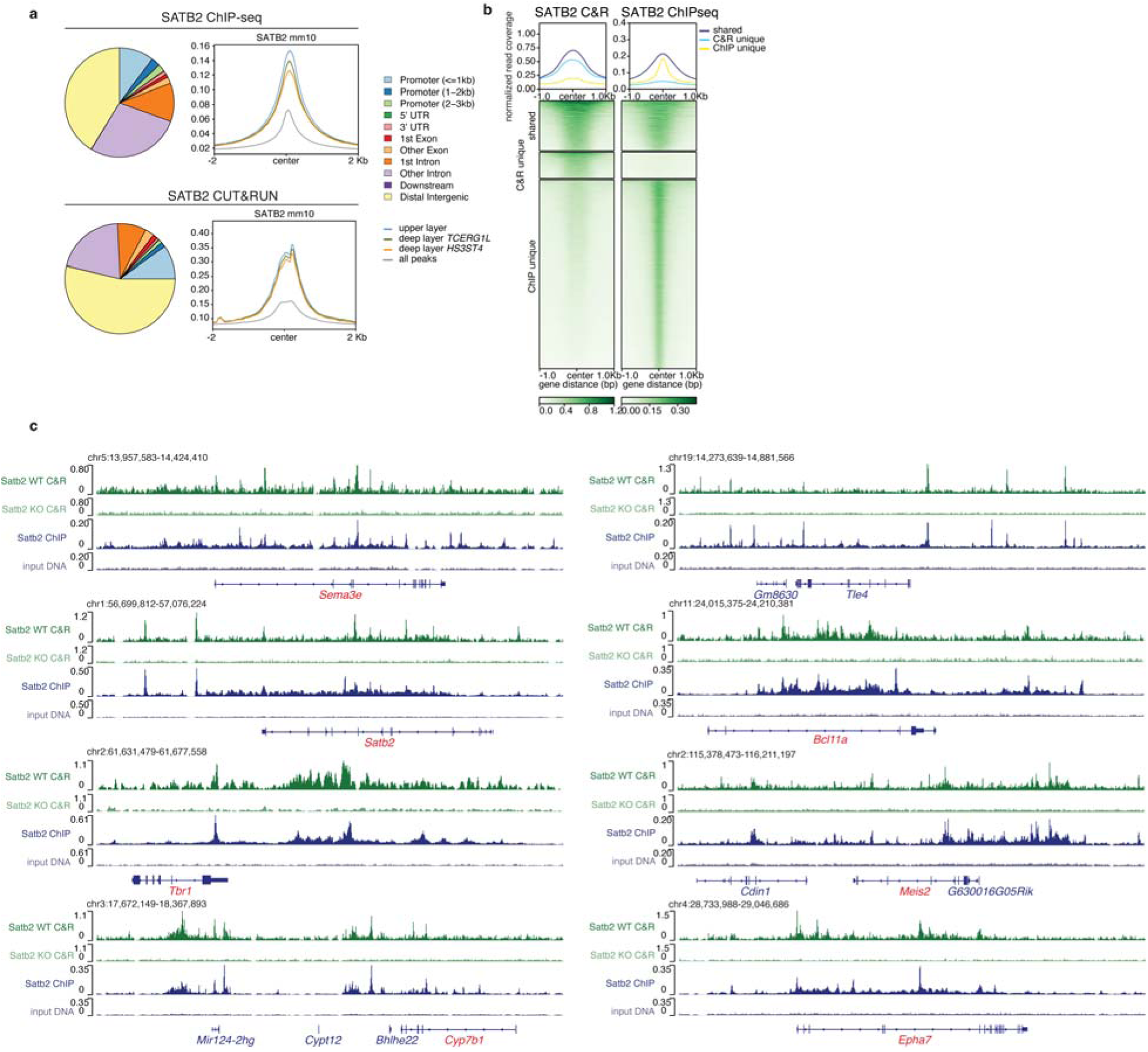
Satb2 CUT&RUN and ChIP-seq reveal concordant genomic binding in mice. **a**, Pie charts showing the distribution of SATB2 binding peaks across genomic regions. **b**, Top, normalized read density for SATB2 CUT&RUN and SATB2 ChIP-seq peaks. Bottom, heatmaps illustrating shared peaks, CUT&RUN unique peaks, and ChIP-seq unique peaks across both datasets. **c**, Genomic browser tracks demonstrating SATB2 binding at the *Sema3e*, *Satb2*, *Tbr1*, *Bhlhe22*, *Cyp7b1*, *Tle4*, *Bcl11a*, *Meis2*, and *Epha7* loci by CUT&RUN and ChIP-seq. CUT&RUN data from *Satb2^lacZ/lacZ^* cortices are included as a control. Genes labeled in red are associated with intelligence.

**Extended Data Figure 2:**
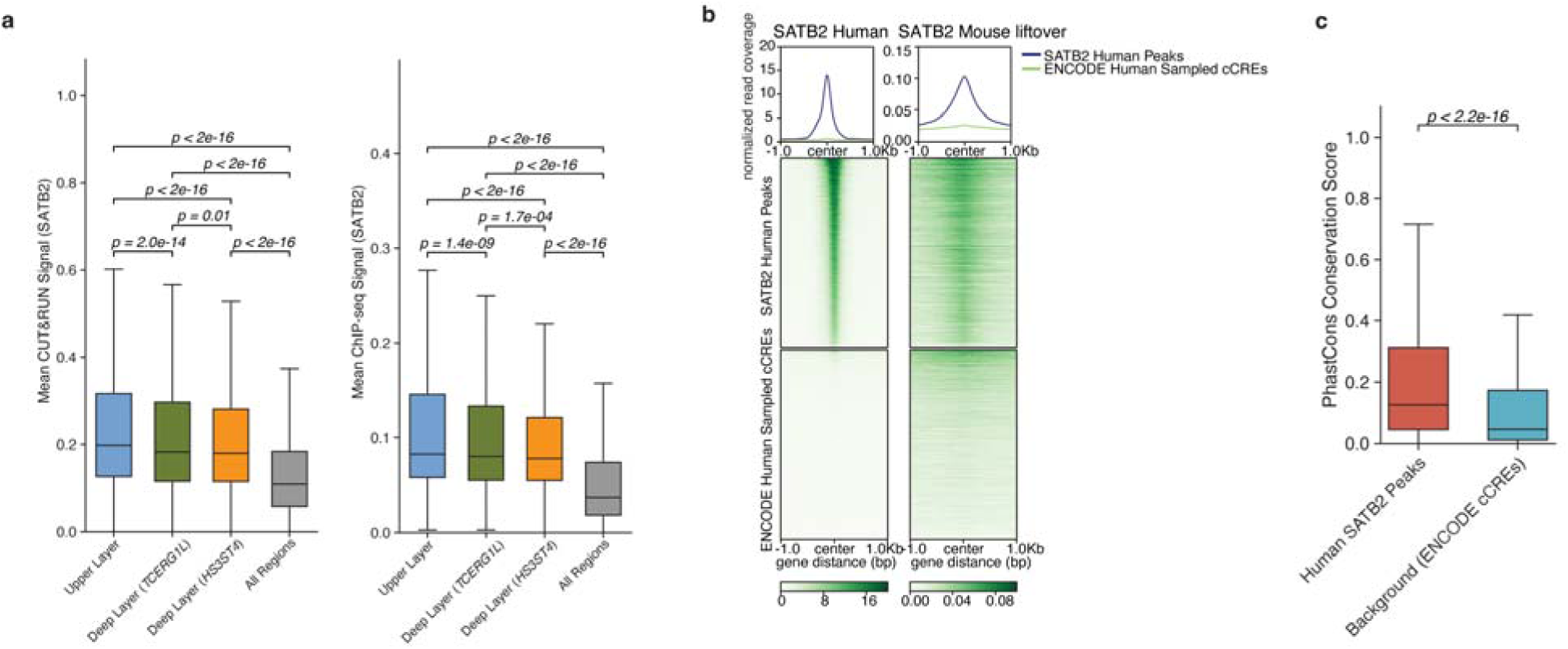
Conservation of SATB2 binding sites in mouse and human genomes. **a**, Enrichment of SATB2 binding in upper-layer, deep-layer TCERG1L^+^ neurons, deep-layer HS3ST4^+^ neurons, and excitatory neurons, based on integrated SATB2 CUT&RUN, SATB2 ChIP-seq, and E18 mouse snMultiome datasets^22^. **b**, Top: Normalized read density for human SATB2 ChIP-seq^25^ and our mouse SATB2 ChIP-seq (lifted over to the human genome). Bottom: Heatmaps showing overlap of human SATB2 peaks and control human cCREs overlapping with human SATB2 ChIP-seq^25^ and our mouse SATB2 ChIP-seq peaks. **c**, Evolutionary conservation scores comparing our mouse SATB2 ChIP-seq peaks with human SATB2 ChIP-seq peaks and background control human cCREs. Statistical significance was assessed using a Wilcoxon two-sided rank-sum test. In box plots, the centre line indicates the median, the box limits denote the 25-75th percentiles, and the whiskers show 1.5x the IQR.

**Extended Data Figure 3:**
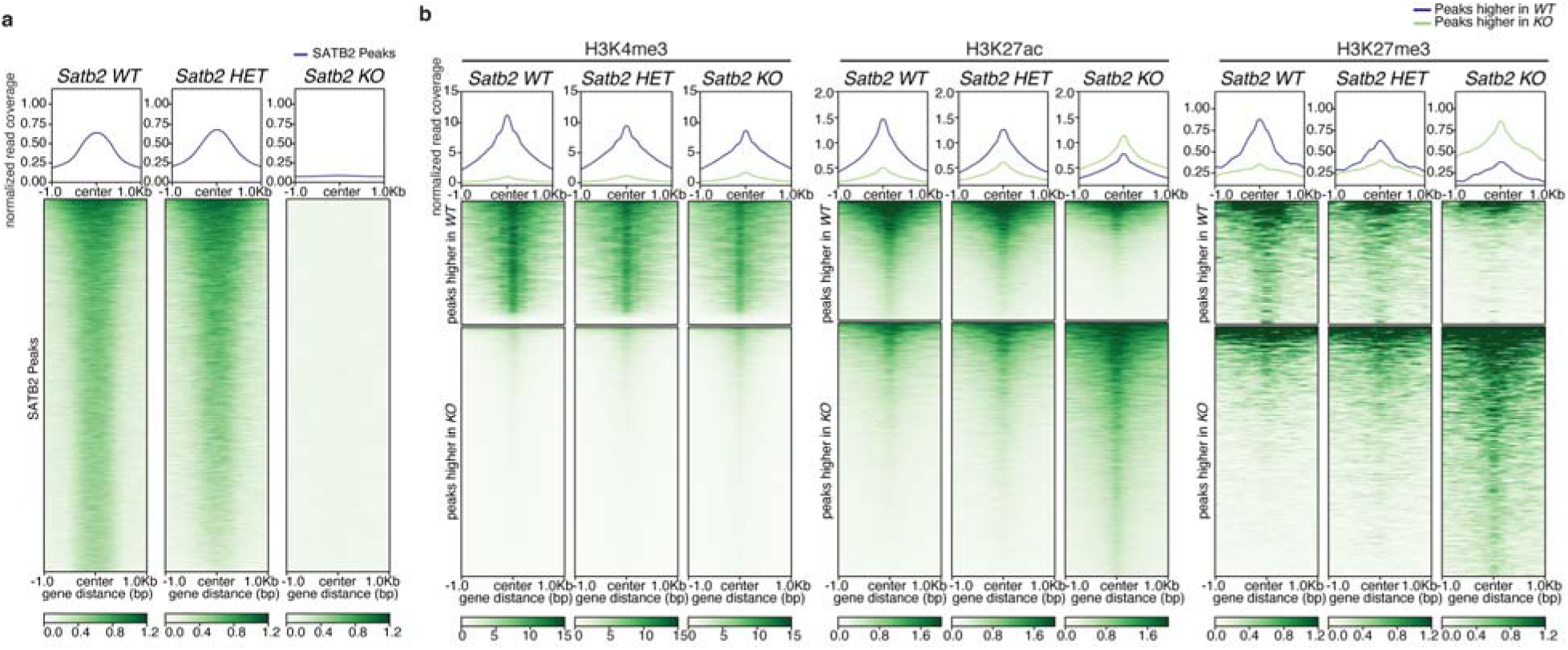
Histone modifications are sensitive to SATB2 dosage. **a**, Top: Normalized read density at SATB2 CUT&RUN peaks in *Satb2 WT, Satb2 HET*, and *Satb2 KO* P0 cortices. Bottom: Heatmaps showing SATB2 CUT&RUN signal across the same peak set in *Satb2 WT, Satb2 HET*, and *Satb2 KO* cortices. **b**, Histone modification CUT&RUN in *Satb2 WT, Satb2 HET*, and *Satb2 KO* cortices. Top: Normalized read density for H3K4me3, H3K27ac, and H3K27me3. Bottom: Heatmaps of differentially bound peaks with significantly higher signal in *WT* cortices (top) or *KO* cortices (bottom), as determined by DiffBind (FDR < 0.10, |log2FC| ≥ 0.26).

**Extended Data Figure 4:**
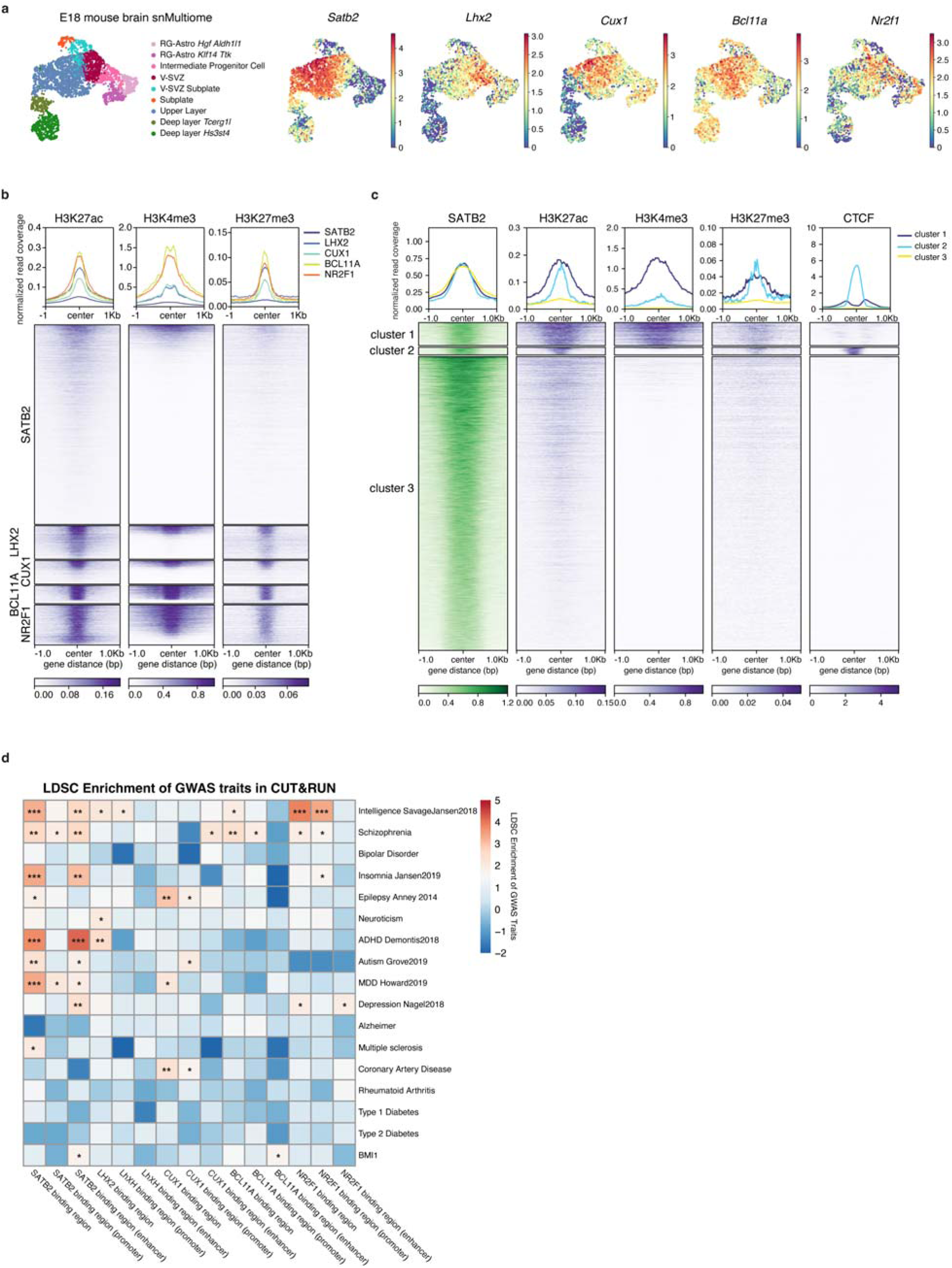
SATB2 CUT&RUN peaks overlap with other transcription factors and are enriched in neurodevelopmental diseases. **a**, Expression UMAP generated from E18 snMultiome data^22^ colored based on cell-type in the first panel, or colored by the normalized expression level of the indicated gene. **b**, Top: Normalized read density of H3K27ac, H3K4me3, and H3K27me3 CUT&RUN peaks from *WT* P0 cortices, showing their overlap. Bottom: Heatmaps of SATB2, LHX2, CUX1, BCL11A, and NR2F1 peaks intersecting histone CUT&RUN peaks. **c**, Top: Normalized read density of regions overlapping CUT&RUN peaks. Bottom: Heatmaps highlighting Cluster 1 (active promoters), Cluster 2 (CTCF binding sites), and Cluster 3 (putative enhancers). **d**,Transcription factor-stratified lineage disequilibrium score regression (LDSC) heatmaps using GWAS summary statistics, integrating multiple transcription factors’ CUT&RUN peaks. \**p* < 0.05; \*\**p* < 0.01; \*\*\**p* < 0.001. Exact *p* values are provided in **Supplementary Table 2**.

**Extended Data Figure 5:**
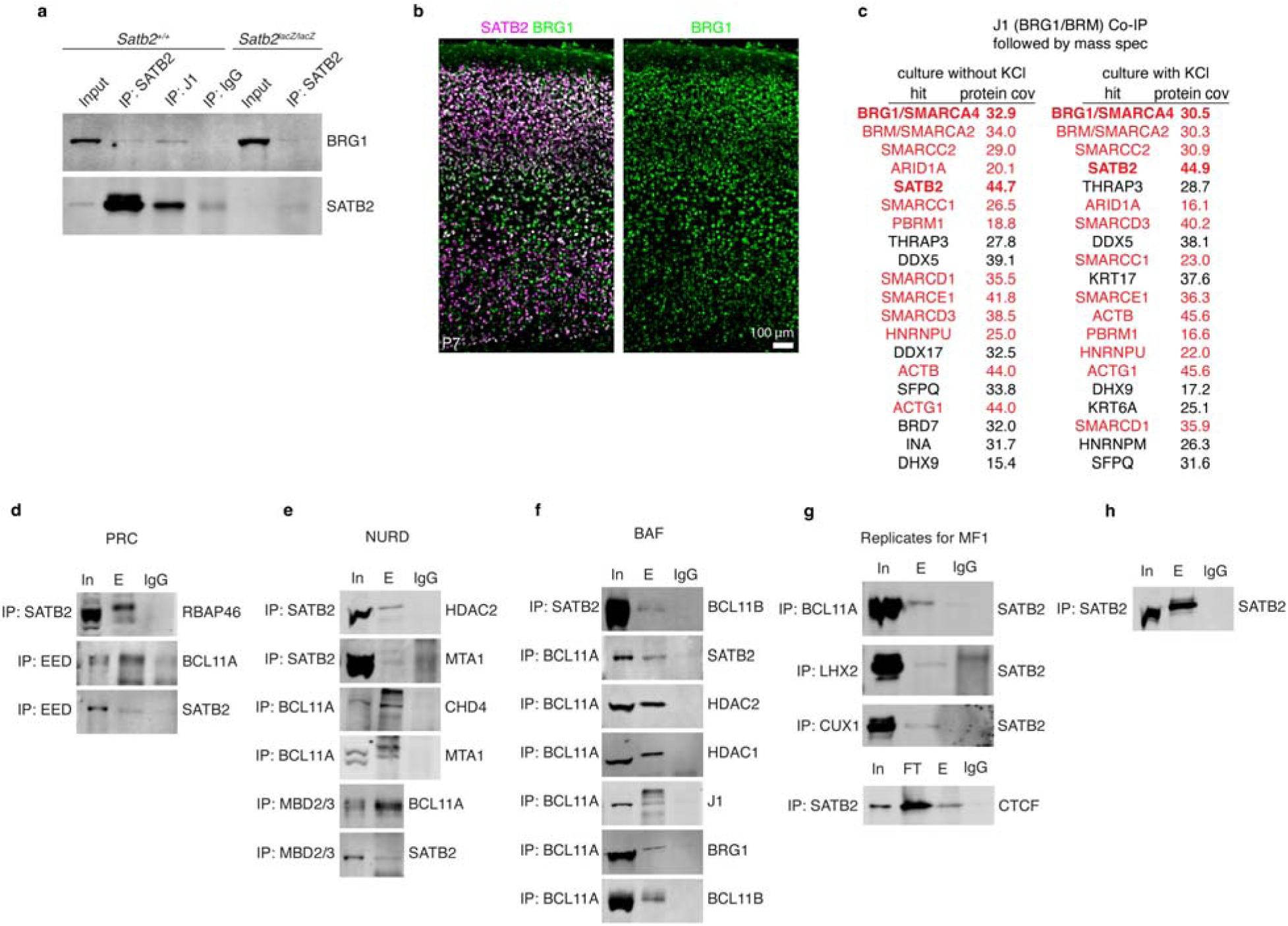
SATB2 cooperates with BAF, NURD, and PRC2 complexes. **a**, Western blot analysis of SATB2 and BRG/SMARCA4 after Co-IP experiments using SATB2, J1, or negative control IgG antibodies and protein extracts from P0 *Satb2^+/+^* or *Satb2^lacZ/lacZ^* cortices. **b**, Immunostaining for SATB2 and BRG1/SMARCA4 in SSp at P7. Scale bar, 100 µm**. c**, The top 20 most enriched proteins immunoprecipitated from cortical neurons using the J1 antibody for BTG1/SMARCA4 and SNF2L2/BRM/SMARCA2^34^. Cortical neurons were cultured with or without KCl in the media. Red text: ID genes. **d-f**, Western blot analysis of PRC (**d**), NURD (**e**), or BAF (**f**) components after Co-IP experiments showing interactions with Satb2 and intact complexes. Protein extracts from P0 *Satb2^+/+^* cortices were used. **g**, Western blots of Co-IP replicates for Fig. 1 showing SATB2 interacting with BCL11A, LHX2, CUX1, and CTCF. Different P0 *Satb2^+/+^* cortices were used than in Fig.1. **h**, Western blot of SATB2 antibody Co-IP validation.

**Extended Data Figure 6:**
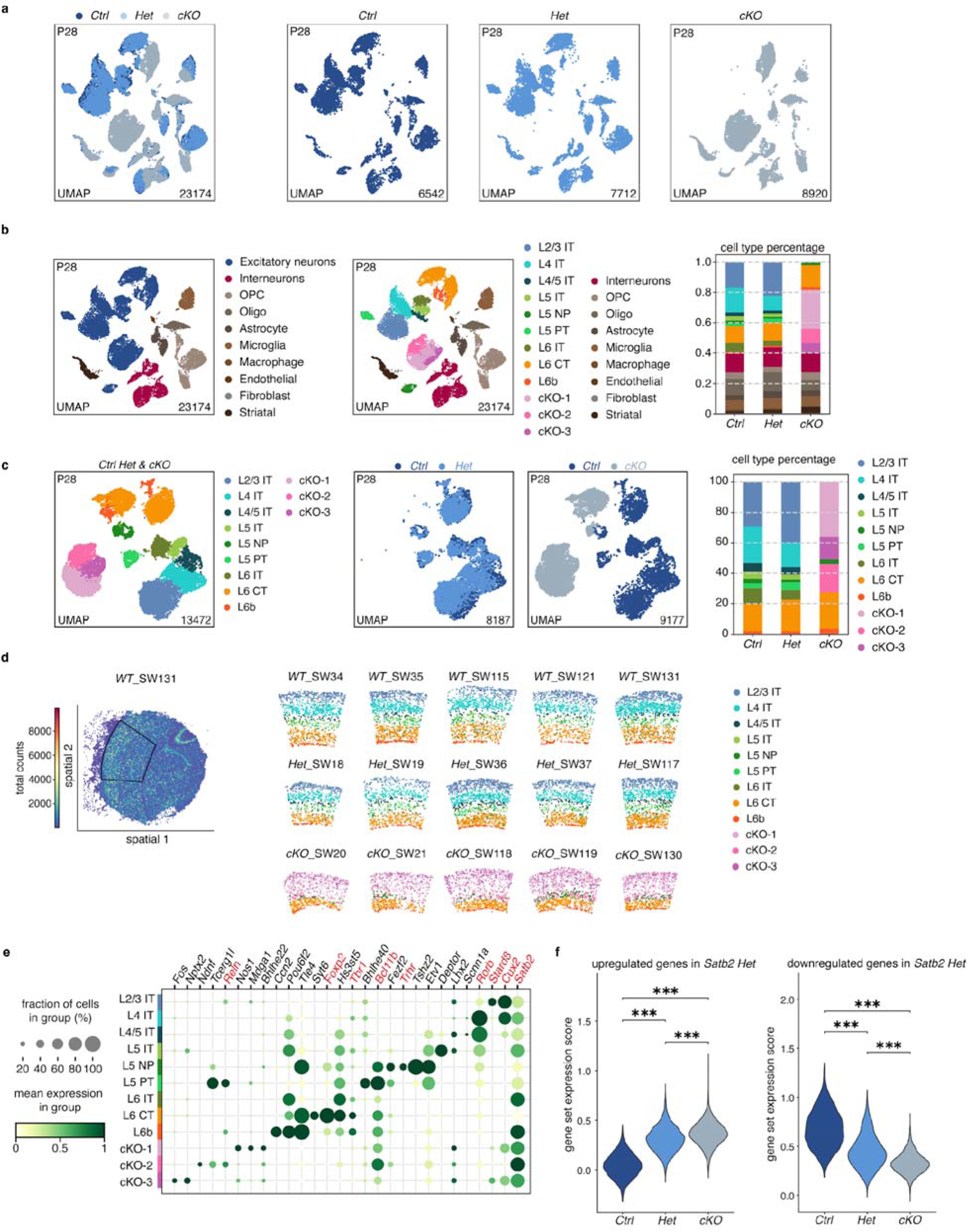
SnRNA-seq analysis for cells in the SSp of control, *Satb2 Het* and *Satb2 cKO* cortices. **a**, UMAP illustrating *Ctrl*, *Het*, and *cKO* clusters overlapped, left, and individual UMAPs for *Ctrl*, *Het*, and *cKO* cells, right. Numbers in the right corner of each UMAP showing the numbers of cells in the samples. (*n* = 2 brains per genotype, 1 female and one 1 male per genotype, single batch, littermates). **b**, UMAPs of all cells from snRNA-seq of P28 *Satb2^+/fl^* (*Ctrl*), *Satb2^fl/lacZ^* (*Het*), and *Satb2^fl/lacZ^;Emx1^Cre^* (*cKO*) primary somatosensory cortex regions. Right, all cell types shown as a percent of total cells identified in *Ctrl*, *Het*, *cKO* snRNA-seq datasets. (*n* = 2 brains per genotype, 1 female and one 1 male per genotype, single batch, littermates). **c**, UMAPs of excitatory neuron clusters in *Satb2^+/fl^* (*Ctrl*), *Satb2^fl/lacZ^*(*Het*), and *Satb2^fl/lacZ^; Emx^cre/+^* (*cKO*) overlaid, colored according to excitatory neuron subtype and mouse genotype. Right, bar graph showing the portions of excitatory neurons belonging to identified excitatory neuron clusters in *Ctrl*, *Het*, *cKO* snRNA-seq datasets. (*n* = 2 brains per genotype, 1 female and one 1 male per genotype, single batch, littermates). **d**, Representative Slide-seq V2 images of *Ctrl*, *Het*, and *cKO* SSp regions labeled by excitatory neuron subtypes. **e**, Dot plot illustrating the top marker genes in each excitatory neuron subtype cluster. The mean expression per group and fraction of positive cells per group are illustrated by color and dot size, respectively. Red text: ID genes. **f**, Violin plots of all upregulated (left) or downregulated (right) DEGs in *Het* cortices compared to *Ctrl*, and *cKO* cortices (Wilcoxon two-sided rank-sum test without correction for multiple comparisons, all *p < 2.22e-16*).

**Extended Data Figure 7:**
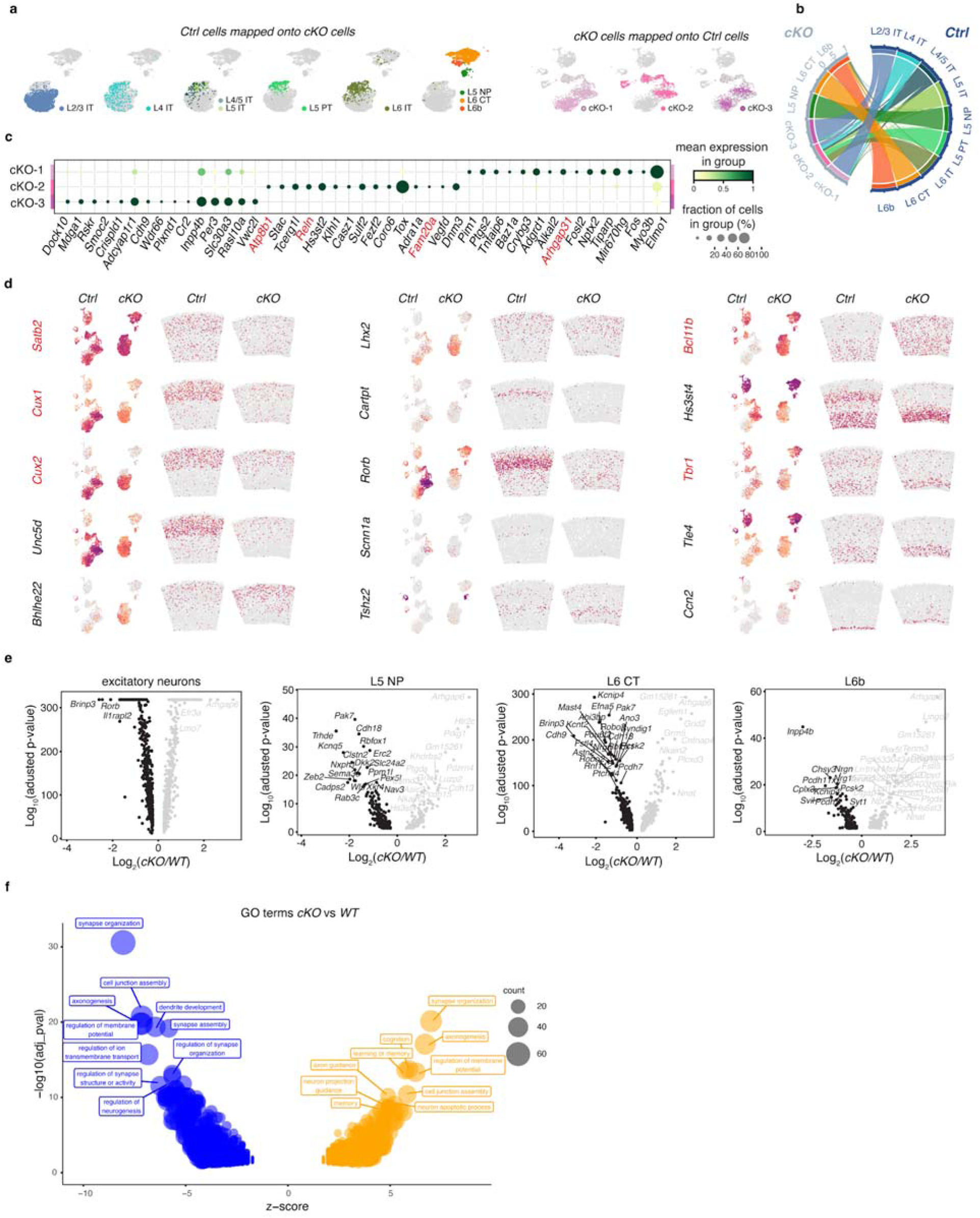
SnRNA-seq and Slide-seq V2 analyses revealed transcriptomic and cell identity changes in the *Satb2 cKO* cortices. **a**, Left, UMAPs of excitatory neuron clusters in *Ctrl* sample mapped onto *Satb2 cKO* UMAPs, colored according to cell type. Right, UMAPs of excitatory neuron clusters in *cKO* mapped onto *Ctrl* UMAPs, colored according to cell type. **b**, Chord diagram illustrating the transcriptomic correspondence between *Ctrl* and *cKO* cortical excitatory neuron clusters determined using TACCO. **c**, Dot plot showing top 15 marker genes for the 3 excitatory neuron clusters, cKO-1, cKO-2, and cKO-3, unique to *Satb2 cKO*. The mean expression per group and fraction of positive cells per group are illustrated by color and dot size, respectively. Red text: ID genes. **d**, UMAPs and Slide-seq V2 signal intensity for known excitatory neuron subtype marker genes. Red text: ID genes. **e**, Volcano plots of DEGs in *cKO* versus *Ctrl* cortices from all excitatory neurons, L5 NP, L6 CT, and L6b clusters. **f**, Gene ontology analysis of all DEGs in *Satb2 cKO* cortices versus *Ctrl* cortices, showing their z-score and unadjusted *p* values. Size of bubbles represents the number of DEGs associated with that term and the z-score represents the ratio of upregulated versus downregulated genes in the *Het* or *cKO* compared to *Ctrl*. Text boxes indicate the top 10 GO terms for up– or downregulated genes.

**Extended Data Figure 8:**
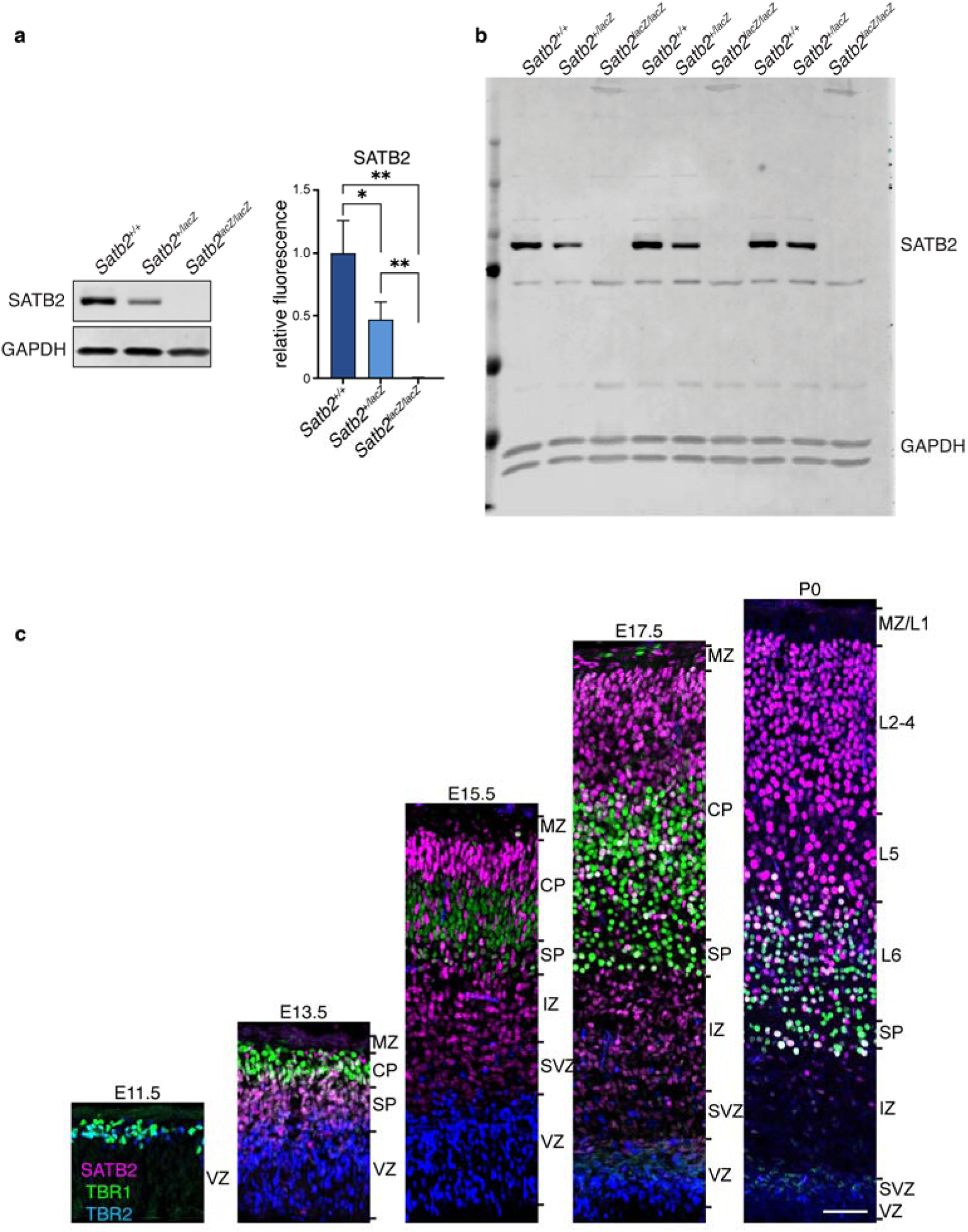
*Satb2^+/lacZ^* mice show decreased protein production. **a**, Western blot for SATB2 using protein extracts from P0 *Satb2^+/+^*, *Satb2^+/lacZ^*, and *Satb2^lacZ/lacZ^* cortices. Signal intensities were normalized to a GAPDH protein loading control. (Unpaired *t*-test; *n* = 3 samples per brain and *n* = 3 brains per genotype; *Satb2^+/+^* vs *Satb2^+/lacZ^ p = 0.0359*; *Satb2^+/+^* vs *Satb2^lacZ/lacZ^ p = 0.0027*; *Satb2^+/lacZ^* vs *Satb2^lacZ/lacZ^ p = 0.0045*.) **b**, Unedited Western blot showing replicates used for analysis stained for SATB2 and GAPDH loading control. **c**, Immunostaining of SATB2, TBR1, and TBR2 on wild type brains from E11.5 to P0. Scale bar 100 µm.

**Extended Data Figure 9:**
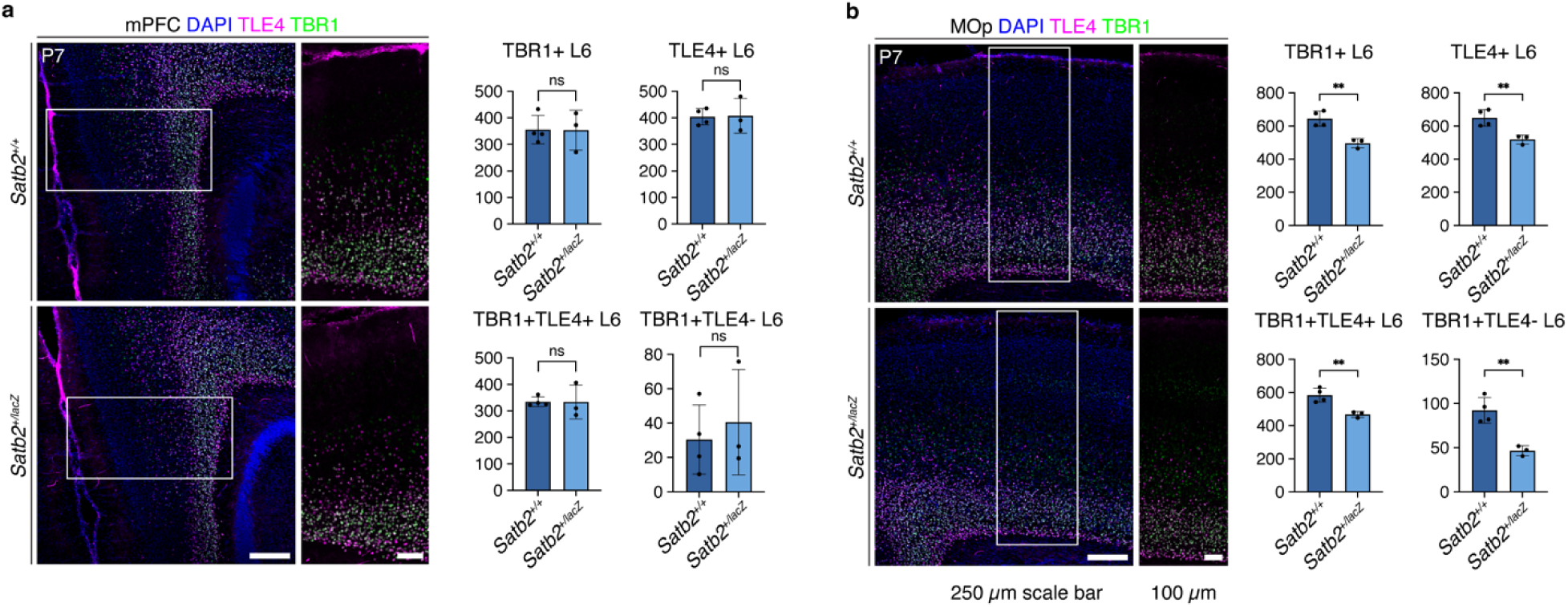
Reduced SATB2 results in laminar defects in MOp but not in mPFC. **a**, Immunostaining of the mPFC in P7 *Satb2^+/+^* and *Satb2^+/lacZ^* brains with TLE4 and TBR1 antibodies. Quantifications of L6 TBR1^+^, TLE4^+^, TBR1^+^ TLE4^+^, and TBR1^+^ TLE4^-^ populations. *n* = 4 *Satb2^+/+^*, *n* = 3 *Satb2^+/lacZ^*. TBR1^+^ L6, *p* = 0.9729; TLE4^+^ L6, *p* = 0.9169; TBR1^+^TLE4^+^ L6, *p* = 0.9817; TBR1^+^TLE4^-^ L6, *p* = 0.6156; unpaired *t*-test. **b**, Immunostaining of the MOp in P7 *Satb2^+/+^* and *Satb2^+/lacZ^*brains with TLE4 and TBR1 antibodides. Quantifications of L6 TBR1^+^, TLE4^+^, TBR1^+^ TLE4^+^, and TBR1^+^ TLE4^-^ populations. *n* = 4 *Satb2^+/+^*, *n* = 3 *Satb2^+/lacZ^*. TBR1^+^ L6, *p* = 0.0044; TLE4^+^ L6, *p* = 0.0096; TBR1^+^TLE4^+^ L6, *p* = 0.0067; TBR1^+^TLE4^-^ L6, *p* = 0.0038. For all graphs: error bars represent ± standard deviation (SD), ns, not significant; \**p* < 0.05, \*\**p* < 0.01, \*\*\**p* < 0.001.

**Extended Data Figure 10:**
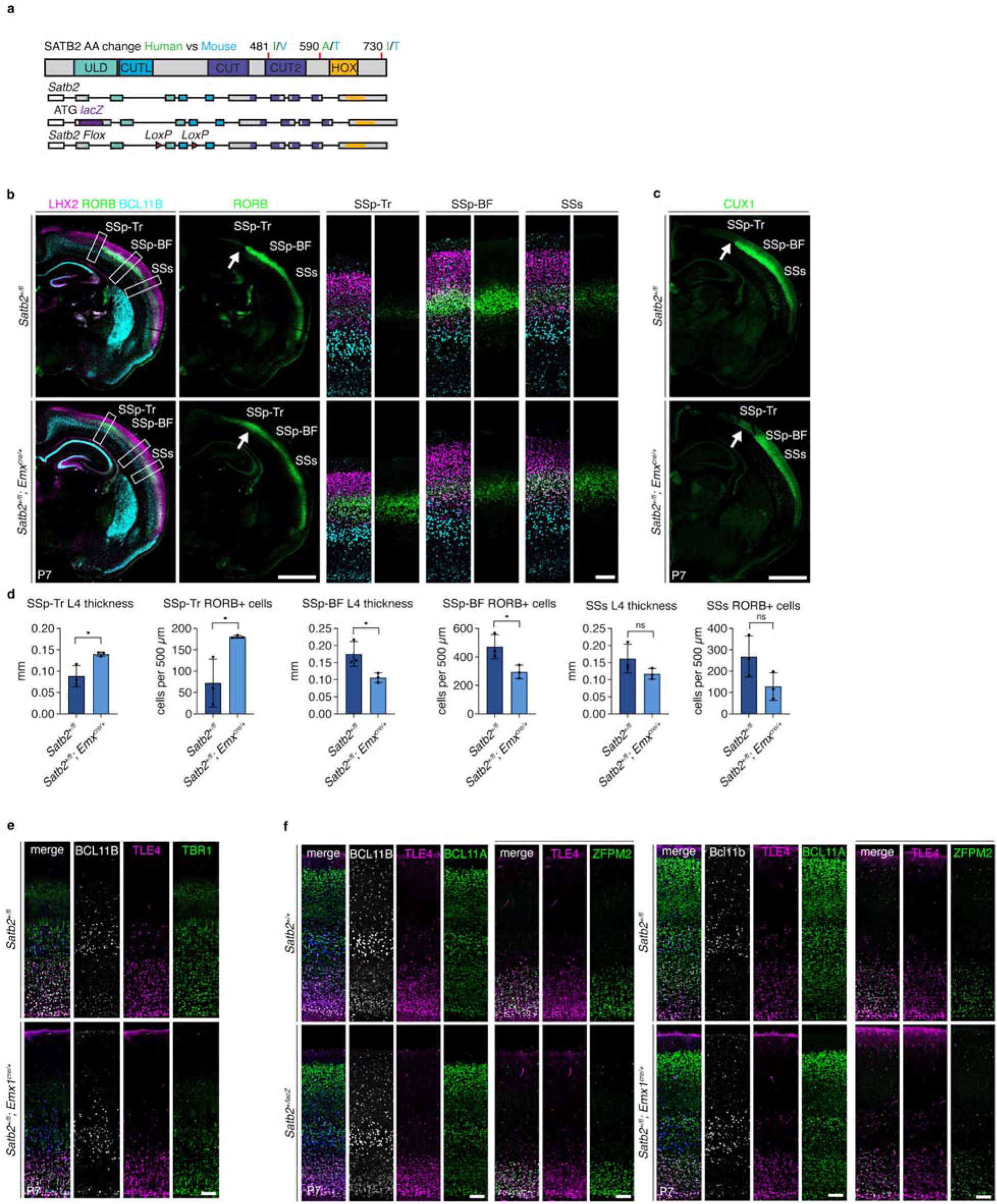
Reduced SATB2 results in decreased L4 thickness and patterning defects, consistent across mutant alleles. **a**, Amino acid differences between mouse and human SATB2 proteins, and *lacZ* and *LoxP* sites in the *Satb2* alleles used in this study. **b**, Immunostaining for LHX2, RORB, and BCL11B in P7 *Satb2^+/+^* and *Satb2^+/fl^*; Emx^cre/+^ brains. White arrows highlight the SSp-Tr regions. SSp-Tr, primary somatosensory area, trunk; SSp-BF, primary somatosensory area, barrel field; SSs, secondary somatosensory cortex. **c**, Immunostaining for CUX1 in P7 *Satb2^+/+^*and *Satb2^+/fl^*; Emx^cre/+^ brains. White arrow highlights the SSp-Tr region. **d**, Quantifications of L4 thickness and RORB^+^ cells per 500-µm wide brain section of indicated region in P7 *Satb2^+/+^*and *Satb2^+/fl^*; Emx^cre/+^ brains. (*n* = 3 brains per genotype. SSp-Tr L4 thickness, *p = 0.024*; SSp-Tr Rorb+, *p = 0.0291*; SSp-BF L4 thickness, *p = 0.0367*; SSp-BF Rorb^+^, *p = 0.034*; SSs L4 thickness, *p = 0.1578*; SSs Rorb^+^ cells, *p = 0.1136*, unpaired *t*-tests. For all graphs: error bars represent ± SD, ns, not significant; \**p* < 0.05.) **e**, Immunostaining of P7 *Satb2^+/fl^* and *Satb2^+/fl^; Emx^cre/+^* conditional het brains with BCL11B, TLE4, and TBR1 antibodies. Scale bar 100 µm. **f**, Immunostaining of BCL11B, BCL11A, TLE4, and ZFPM2 in P7 *Satb2^+/+^* and *Satb2^+/lacZ^*brains (left) or *Satb2^+/fl^* and *Satb2^+/fl^; Emx^cre/+^* conditional het brains (right). Scale bar 100 µm.

**Extended Data Figure 11:**
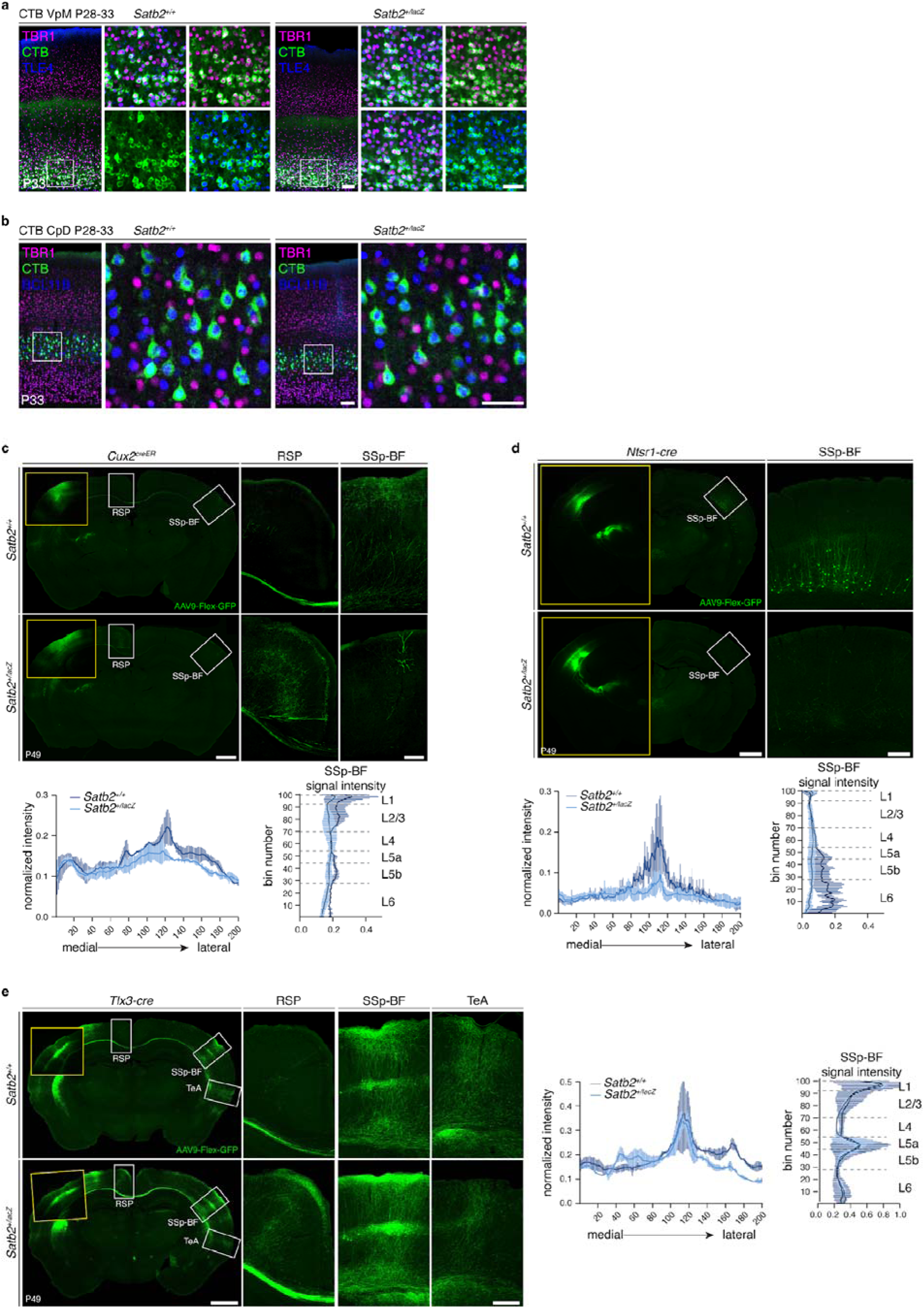
*Satb2^+/lacZ^* mice show altered IT projections. **a-b**, CTB injections into the ventral posteromedial nucleus (VpM) (**a**) and cerebral peduncle (CpD) (**b**) label L6 CT and L5 PT neurons in P33 *Satb2^+/+^*and *Satb2^+/lacZ^* brains. TBR1 and TLE4 (**a**), and TBR1 and BCL11B (**b**) immunostainings are shown. White boxes indicate magnified areas in L6 or L5 in the primary somatosensory cortex contralateral to the injection side. Scale bars: low magnification, 1 mm; high magnification, 100 µm. **c-e**, Images and quantifications of GFP fluorescence signal in *Satb2^+/+^; Cux2-Cre^ER^*and *Satb2^+/lacZ^; Cux2-Cre^ER^* **(c)**, *Satb2^+/+^; Ntsr1^cre/+^* and *Satb2^+/lacZ^; Ntsr1^cre/+^***(d)**, and *Satb2^+/+^; Tlx3^cre/+^* and *Satb2^+/lacZ^; Tlx3^cre/+^* **(e)** brains which were injected with AAV9-FLEX-GFP virus in SSp at P28. Quantifications show the GFP fluorescence signal along medial-lateral axis of the contralateral cortical plate (left), and normalized GFP fluorescence along the apical-basal axis of the contralateral SSp-BF. Scale bars: low magnification, 1 mm; high magnification, 100 µm.

**ExDF 12:**
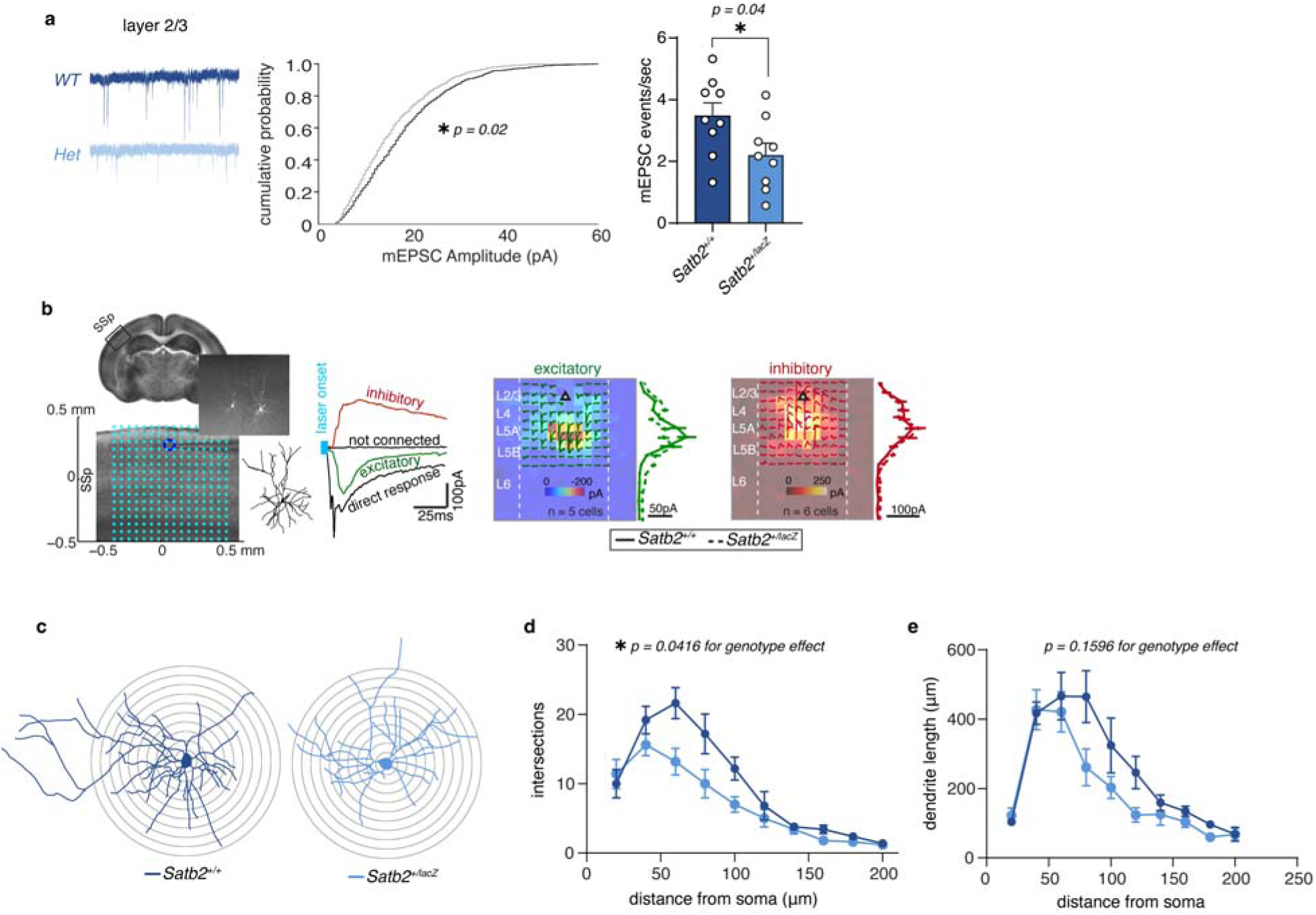
Electrophysiological changes of L2/3 IT neurons in *Satb2^+/lacZ^* mice. **a**, Representative mEPSC responses from *Satb2^+/+^* and *Satb2^+/lacZ^* L2/3 IT neurons. Cumulative distribution of pooled mEPSCs, and frequency of events, (*n* = 9 *Ctrl* and 9 *Het*, *p = 0.02* on cumulative distribution, K-S test, and *p = 0.04* on frequency of mEPSCs, two-tailed Mann-Whitney U test). **b**, Diagram illustrating the SSp region and laser-scanning grid in LSPS with glutamate uncaging experiments for L2/3 IT neurons. Example responses of excitatory, inhibitory, and direct response signals. Representative LSPS maps and pooled EPSC (green) or IPSC (red) responses. Color maps represent averaged responses from 5-6 *Satb2^+/+^* neurons and averaged synaptic responses (mean ± SD) are binned by laminar location and presented to the right of the colormap. **c**, Representative Sholl profiles of L2/3 IT neurons from *Satb2^+/+^* and *Satb2^+/lacZ^*cortices. **d**, Sholl intersection profiles of L2/3 IT *Satb2^+/+^*and *Satb2^+/lacZ^* neurons (*n* = 5 *WT* and *Het* neurons, Genotype effect: F_(1,8)_ = 5.875, *p = 0.0416*, two-way ANOVA). **e**, Dendrite length profiles of L2/3 IT neurons from *Satb2^+/+^*and *Satb2^+/lacZ^* cortices (*n* = 5 *WT* and *Het* neurons, Genotype effect: F_(1,8)_ = 2.404, *p = 0.1596*, two-way ANOVA).

**Extended Data Figure 13:**
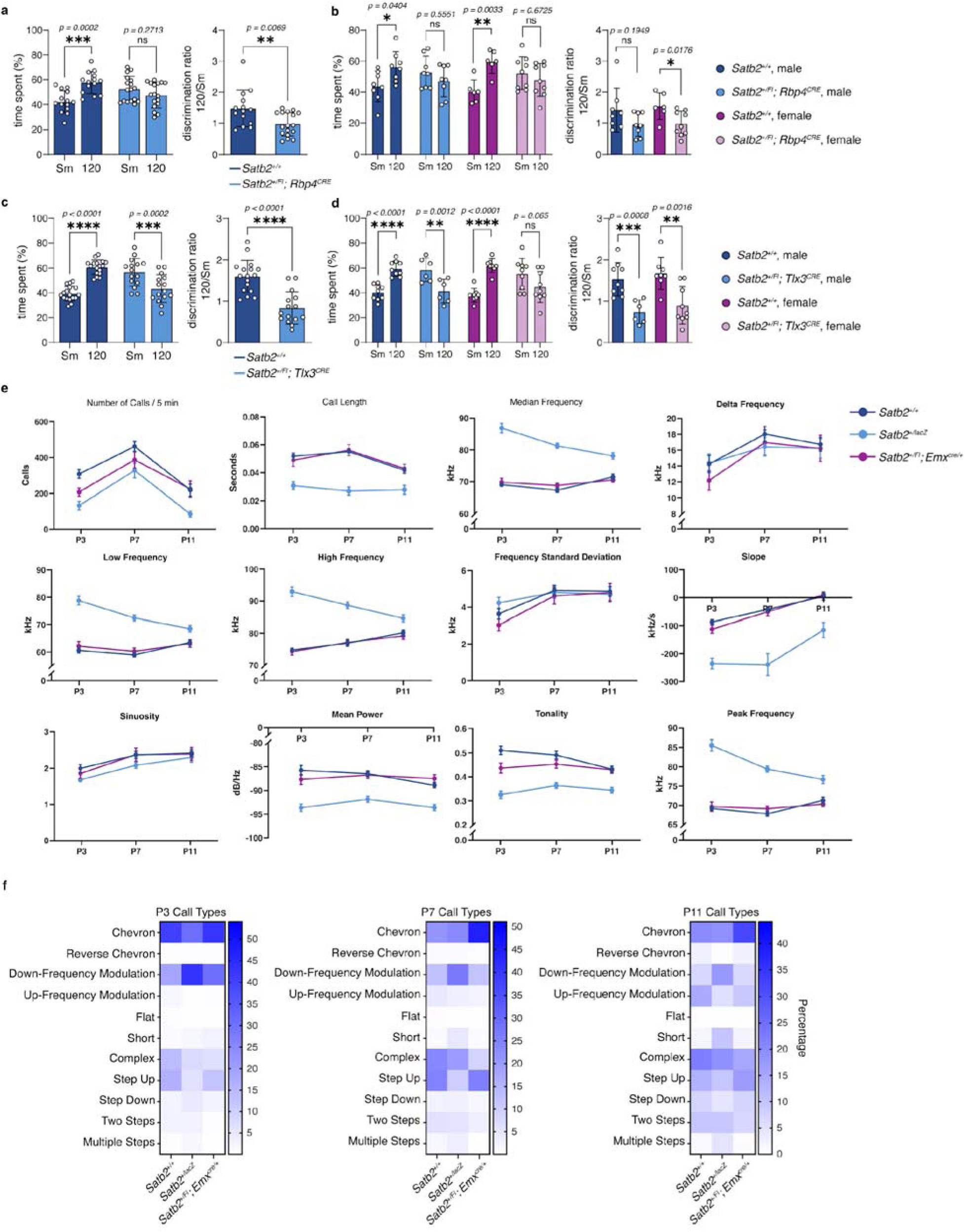
Layer-specific reduction of SATB2 leads to for defective sensory discrimination, and USVs are not dependent on cortical SATB2. **a**, *Satb2^+/+^* vs *Satb2^+/fl^; Rbp4^cre/+^*texture discrimination between smooth (Sm) or 120-grit (120) sandpaper quantified as percentage of time spent per texture and the discrimination ratio (*n* = 14 *WT* and 17 *Rbp4*-*Het*, Two-Way Anova with Bonferroni correction; discrimination ratio: Mann-Whitney test). **b**, Texture discrimination from **a** separated by sex and genotype. Percentage of time spent per texture and the discrimination ratio are shown. (*n* = 8 *WT* male, 6 *WT* female, 8 *Rbp4*-*Het* male, and 9 *Rbp4*-*Het* female, time spent, Two-Way Anova with Bonferroni correction; discrimination ratio, Mann-Whitney test, error bars represent ± SEM). **c**, *Satb2^+/+^* vs *Satb2^+/fl^; Tlx3^cre/+^* texture discrimination between smooth (Sm) or 120-grit (120) sandpaper quantified as percentage of time spent per texture and the discrimination ratio (*n* = 17 *WT* and 15 *Tlx3-Het*, Two-Way Anova with Bonferroni correction; discrimination ratio: Mann-Whitney test). **d**, Texture discrimination from **c** separated by sex and genotype. Percentage of time spent per texture and the discrimination ratio are shown. (*n* = 9 *WT* male, 8 *WT* female, 6 *Tlx3-Het* male, and 9 *Tlx3-Het* female, time spent, Two-Way Anova with Bonferroni correction; discrimination ratio, Mann-Whitney test, error bars represent ± SEM). **e**, Timeline of isolation induced ultrasonic vocalization acoustic analysis between *Satb2^+/+^*, *Satb2^+/lacZ^* and *Satb2^+/fl^; Emx^cre/+^* mice at P3, P7, and P11 (Detailed statistics in **Supplementary Table 1**). **f**, Heatmaps of call type percentages from *Satb2^+/+^*, *Satb2^+/lacZ^*and *Satb2^+/fl^; Emx^cre/+^* mice at P3, P7, and P11.

## Mice used in this study

Experiments were performed according to protocols approved by the Institutional Animal Care and Use Committee at University of California at Santa Cruz, at University of Arizona College of Medicine Phoenix, and at Stanford University. Experiments were performed in accordance with institutional and federal guidelines.

The day of the vaginal plug detection was designated as E0.5. The day of birth was designated as P0. The sexes of the embryonic and early postnatal mice were not determined. The *Satb2^+/lacZ^*^53^, *Satb2^flox^*^/+ 54^, *Emx1^Cre^*^55^, *Tlx3^cre^*^/+ 56^, *Cux2^creER^*^/+ 57^, *Rbp4^cre^*^/+ 58^, and *Ntsr1^cre^*^/+ 58^ mice were described previously. The following mice were used in this study: *Satb2*^+/+^ and *Satb2^+/lacZ^* mice: P0, P1, P4, P7, P28, P30, and adult (≥ P60), both male and female mice were used. *Satb2^flox/+^,Satb2^flox/lacZ^, Satb2^flox/lacZ^;Emx1^cre/+^ mice: P0, P7, P28 and adult (*≥ *P60), both male and female mice were used. Tlx3^cre/+^*, *Cux2^creER/+^*, and *Ntsr1^cre/+^* mice were used for breeding to *Satb2^+/lacZ^*mice to generate *Satb2^+/lacZ^*; *Tlx3^cre/+^*, *Satb2^+/lacZ^*; *Cux2^creER/+^*, and *Satb2^+/lacZ^*; *Ntsr1^cre/+^* mice which were used at P28 for axon tracing, both male and female.

## Method Details

### Immunohistochemistry

To prepare tissue for staining, mice were anesthetized and prepared for trans-cardiac perfusion. 1X PBS was delivered into the body from the left ventricle. When the fluid exiting the right atrium ran clear, 4% paraformaldehyde was pumped until the body stiffened. Brains were then dissected and post-fixed in 4% paraformaldehyde, 0.1% saponin, 1X PBS overnight at 4°C and then cryoprotected in 30% sucrose 1X PBS, or until the brains sank. The brains were sectioned using a sliding microtome into 50µm sections. Sections were permeabilized with 0.6% Triton X-100 in 1X PBS for 30 minutes before incubating in blocking buffer (5% horse serum, 0.03% Triton X-100, 1X PBS) for 1 hour. Blocking buffer was removed, and the sections were incubated with primary antibodies diluted in blocking buffer overnight at 4°C. The following antibodies were used for immunohistochemistry in this study: SATB2, 1:1000 (rabbit, Abcam, ab34735), TLE4, 1:100 (mouse, Santa Cruz Biotechnology, sc-365406), TBR1, 1:1000 (rabbit, Abcam, ab31940), BRG1, 1:500 (mouse, Santa Cruz Biotechnology, sc-17796), NTNG1 1:100 (goat, R&D Systems, AF1166), PKCG, 1:500 (rabbit, Santa Cruz Biotechnology, sc-211), L1, 1:500 (rat, MilliporeSigma, MAB5272), BCL11B, 1:1000 (rat, Abcam, ab18465), BCL11A, 1:1000 (rabbit, Abcam, ab191401), ZFPM2, 1:1000 (rabbit, Santa Cruz Biotechnology, sc-10755), BHLHE22, 1:100 (goat, Santa Cruz Biotechnology, sc-6045), LHX2, 1:500 (goat, Santa Cruz Biotechnology, sc-19344), RORB, 1:500 (rabbit, Proteintech 17635-1-AP), and CUX1, 1:500 (rabbit, Santa Cruz Biotechnology, sc-6327). The sections were washed three times with 1X PBS 0.03% Triton X-100 for 5 minutes each, and then incubated with secondary antibodies conjugated to Alexa 488, 555, or 647 fluorophores (Jackson), all diluted in blocking buffer at a ratio of 1:1000. Secondary antibodies were incubated for 2 hours at room temperature. Nuclear staining was performed by incubating with 4’,6-Diamidino-2-Phenylindole, Dihydrochloride (DAPI) 1:10,000 (Invitrogen, D1306) in 1X PBS for 10 minutes. Sections were then manually arranged on microscope slides before being coverslipped with Fluoromount-G.

### Cytochrome oxidase staining

Brains used for cytochrome c oxidase staining were dissected from perfused mice as described above. Brains were cut in half along the midline, and all subcortical structures were removed with standard dissecting tools to obtain cerebral cortices. Isolated cortices were flattened between two glass slides with a 1mm spacer in-between and placed in 4%PFA, 0.1% saponin in 1X PBS at 4°C overnight and then cryoprotected by submerging in 30% sucrose, 1X PBS for 24 hours. The flattened cortices were then sectioned on a sliding microtome at 50 µm. Sections were washed with 1X PBS and then incubated at 37°C in CO staining buffer containing: 5% sucrose, 0.03% cytochrome c, 0.02% catalase, 0.05% 3,3′-Diaminobenzidine, and 0.1M phosphate buffer for 2-3 hours. Sections were then washed with 1X PBS and mounted in Fluoromount-G and imaged the following day.

### ChIP-seq

ChIP-seq was performed as described^59^ with modifications. P0 pups were anesthetized on ice and sacrificed. Brains were quickly taken out, neocortices were dissected under a dissection microscope, and the meninges were removed. The tissue was washed in 1X PBS before 2mM DSG in 1X PBS was added. The tissue was immediately triturated into a single-cell suspension, and incubated at room temperature (RT) for 30 minutes. Formaldehyde was added to a final concentration of 1%, and cells were incubated for 10 minutes with rotating at RT. 2M Glycine was added to quench the formaldehyde. The cells were then washed with 1X PBS, resuspended in lysis buffer (1% SDS, 10 mM of EDTA, and 50 mM Tris HCl, pH8.1), and incubated on ice for 10 minutes. Samples were sonicated with a Diagenode Bioruptor 300 for 32 cycles: 30 seconds on, 90 seconds off, at 4°C, and centrifuged at 14,000g for 10 minutes at 4°C. 9 volumes of ice-cold IP dilution buffer (0.01% SDS, 1.1% Triton X-100, 1.2 mM EDTA, 16.7 mM Tris-HCl, pH 8.1, 167 mM NaCl) were added to the supernatant. Sonicated chromatin solution from 20 million cells was used for each ChIP reaction. The mouse monoclonal SATB1/SATB2 antibody (Abcam, ab51502) was used, and 2 biological replicate experiments were performed. 3 µg of antibody was added to each tube and the reactions were incubated overnight at 4°C with gentle mixing. Immunoprecipitation was performed using Protein A and G Dynabeads (NEB, S1425S, S1430S). 20 µl of beads (10 µl Protein A and 10 µl Protein G) were added to each sample and incubated for 2 hours at 4°C. Beads were then washed for 3 minutes each with the following buffers: TSE-150 (1% Triton X-100, 0.1% SDS, 2 mM EDTA, 20 mM Tris-HCl, pH 8.1, 150 mM NaCl), TSE-150 or TSE-500 (1% Triton X-100, 0.1% SDS, 2 mM EDTA, 20 mM Tris-HCl, pH 8.1, 500 mM NaCl), LiCl/detergent wash (0.25 M LiCl,1% NP-40, 1% DOC, 1 mM EDTA, 10 mM Tris-HCl, pH 8.1), and TE (10 mM Tris, pH 8.0, 1 mM EDTA). Samples were then eluted with 250 µl of elution buffer (1% ultrapure SDS, 0.1 M NaHCO_3_), supplemented with NaCl to a final concentration of 0.2 M, and incubated at 65°C overnight to reverse the crosslink.

Samples were treated with RNase A at 37°C for 30 minutes and Proteinase K at 42°C for 2 hours before being extracted Phenol:Chloroform:Isoamyl alcohol using phase-lock tubes (Qiagen 129046). Fragments were then precipitated with 100% EtOH, washed with 75% EtOH, and resuspended in 30 µL water.

### CUT&RUN

CUT&RUN was performed according to published protocol^20^. P0 brains were quickly dissected and dropped into ice cold 1X PBS with protease inhibitor (Roche, 11873580001). The meninges were carefully removed, and the dissected neocortices were cut into small pieces. The tissue was washed with 400 µL ice cold Accumax solution (Sigma-Aldrich A7089). Ice cold Accumax was removed and 400 µL Accumax prewarmed at 37°C was added. The cortical tissue was incubated at 37°C for 5 minutes before being manually dissociated with a P1000 pipet tip. Dissociated cells were spun down at 600 g for 3 minutes and washed with 1 mL ice cold wash buffer (20 mM HEPES pH 7.5, 150 mM NaCl, 0.5 mM Spermidine, 1X EDTA-free protease inhibitor) three times. Cells were resuspended, counted, and aliquoted. 10 µL Concanavalin-A beads (Bangs Laboratories, PB531) were added to each tube containing 500,000 cells. The tubes were placed on a magnetic stand and the wash buffer was removed. 150 µL antibody buffer (wash buffer with 2 mM EDTA, 0.03% Digitonin, MilliporeSigma, 300410) was added to each tube. All antibodies were added at a dilution of 1:100, except the IgG which was added to a final concentration of 1 µg/µL. The following antibodies were used in CUT&RUN experiments: SATB2 (rabbit, Abcam, ab34735), CUX1 (rabbit, Santa Cruz Biotechnology, sc-6327), LHX2 (goat, Santa Cruz Biotechnology, sc-19344), NR2F1 (Novus Biological, NBP1-31259), H3K27me3 (Cell Signaling, 9733S), H3K27Ac (Abcam, Ab4729), H3K4me3 (Cell Signaling, 97515), Rabbit IgG (Invitrogen 02-6102). Samples were incubated at 4°C overnight with shaking at 400 rpm. The following day, beads were washed twice with 1 mL DIG-wash buffer (wash Buffer with 0.03% Digitonin) and resuspended in 50 µL DIG-wash buffer. 2.5 µL A/G-MNase (EpiCypher, 15-1016) was added to 50 µL DIG-wash buffer, and the samples were incubated for 1 hour at 4°C shaking. Beads were washed twice with DIG-wash buffer and resuspended in 100 µL DIG-wash buffer. Ca^2+^ was added to a final concentration of 1 mM and the samples were incubated for 1 hour at 0°C in a heat block immersed in an ice water slurry. The reaction was inhibited by adding 100 µL 2X STOP buffer (170mM NaCl, 20mM EGTA, 0.05%, 50 µg/mL RNase A, 20 µg/mL Glycogen, 250 pg *E. coli* Spike-in DNA, EpiCypher 18-1401). Samples were then incubated at 37°C for 30 minutes, and supernatant was transferred to fresh tubes. Phenol chloroform isoamyl alcohol extraction was performed, and purified CUT&RUN DNA was resuspended in 30 µL 1 mM Tris-HCl pH 8.0, 0.1 mM EDTA.

### ChIP-Seq and CUT&RUN library prep and sequencing

We used the NEBNext Ultra II DNA Library Prep Kit (E7645L) with some modifications to the protocol. For the adapter ligation step, the NEBNext Adaptor for Illumina was diluted 1:50, to 0.3µM. The adaptor ligated DNA was then cleaned up with 2.1X AMPure XP beads (Beckman A63881) before proceeding. PCR was performed with NEBNext Multiplex Oligos for Illumina (E7600S) with the following thermocycler conditions: Initial denaturation 98°C, 30s, denaturation 98°C, 10s, annealing and extension 65°C, 12s, repeat denaturing and annealing and extension 14 times, final extension 65°C, 5 minutes in a Bio-Rad T100 Thermal Cycler. After amplification, 0.5X AMPure beads were added to the reaction, mixed, incubated for 10 minutes, and samples were placed on a magnet stand. Supernatant was transferred to fresh tubes and then cleaned up with 2X AMPure beads. DNA concentrations were determined using Bioanalyzer High Sensitivity DNA kit (5067-4626). DNA between 146 and 600 bp was quantified for the analysis. Samples were pooled such that equimolar amounts of each sample were present in the pool. The primer-dimer peak was removed with Pippin size selection. The pool was sequenced on the NextSeq platform (high output, 75 cycles). The sequencing was demultiplexed, and sequencing reads were mapped to the genome. Binding peaks were called using SEACR^19^.

### Analysis of ChIP-seq and CUT&RUN data

ChIPseeker v1.30.3^60^ was used to annotate DNA binding regions in the Satb2 ChIP-seq and Satb2 CUT&RUN data. We used the computeMatrix, plotHeatmap and plotProfile functions from the deepTools v3.5.1 toolkit to visualize the enrichment patterns of DNA binding and histone states across different peak sets. We used liftOver with ‘-minMatch=0.5’ to lift the mouse Satb2 ChIP-seq data (mm10) to human genome (hg38) to enable comparable visualization with human ATAC-seq data, RNA-seq data and the Proximity Ligation-Assisted H3K4me3 ChIP-seq (PLAC-seq) data^26^. Differential binding between genotypes was assessed using DiffBind (v3.10), with significance thresholds of FDR < 0.10 and |log2FC| ≥ 0.26. To quantify the concordance between the two methods, peak-level overlap between ChIP-seq and CUT&RUN peak sets was assessed using a permutation test (*n* = 1,000 iterations) implemented in regioneR, separately for promoter and distal intergenic peaks.

### snRNA-seq

P28 mice were anesthetized, and brains were quickly dissected and placed into ice cold 1X PBS. The brains were then sectioned at 500 µm using a vibratome. SSp regions were manually excised from the brain sections using small iris scissors and flash frozen in liquid nitrogen. Frozen tissue was stored at –80 °C until further processing. Nuclei were isolated as previously described^61^. Briefly, frozen tissue was immediately immersed in ice-cold Homogenization Buffer (HB) containing 0.25 M sucrose, 25 mM KCl, 5 mM MgCl_2_, 20 mM Tricine-KOH, 1 mM DTT, 0.15 mM spermine, and 0.5 mM spermidine, 0.25% IGEPAL-CA630, and 0.2 U/µL RNasin. Tissue homogenization was performed with 8-10 strokes of pestle A, followed by a 5 min wait, and then 8-10 strokes of pestle B using a 2 mL Dounce tissue grinder. Following centrifugation and resuspension, DRAQ5 (BioLegend) was added to the nuclei suspension for sorting on a Sony MA900 cell sorter using a 70 µm nozzle. Nuclei were collected in pre-chilled 0.2 ml PCR tube and counted using a hemocytometer (INCYTO C-Chip). snRNA-seq libraries were prepared with Chromium Single Cell 3’ Kit v3.1 (10x Genomics) according to manufacturer’s protocol. Pooled libraries were sequenced on NovaSeq 6000 instruments (Illumina).

### Slide-seq V2 library preparation

Slide-seq pucks (round, 3 mm in diameter) were generated as described previously^62^. 10 µm-thick coronal sections were obtained from flash-frozen brain samples using cryostat (Leica) and used for generating Slide-seq library immediately, following published Slide-seqV2 protocol^36^ (also on protocol.io https://www.protocols.io/view/library-generation-using-slide-seqv2-bxijpkcn). Libraries were pooled and sequenced on NovaSeq 6000 flow cells (Illumina).

### snRNA-seq data processing

CellRanger (v7.0.0, 10x Genomics) was used with default parameters to map snRNA-seq data to the mouse reference genome (mm10) provided by 10x Genomics. The gene expression matrices output from CellRanger were then imported into Python v3.9.13 as AnnData objects (anndata v0.8.0). Scanpy v1.9.1 was used as the basic framework for snRNAseq processing. We used Scrublet v.0.2.3^63^ to calculate the potential doublet score for each cell. Unless specified elsewhere, genes expressed in fewer than 3 cells were filtered. Cells were filtered based on the following criteria: n_counts□<□15,000, 200□<□n_genes□<□4,000, ratio_of_mitochondria_genes□<□5% (gene symbols beginning with ‘mt-’), ratio_of_ribosome_genes□<□5% (gene symbols beginning with ‘Rpl’ or ‘Rps’) and scrublet_score < 0.25. After quality control, the raw counts were normalized using the pp.normalize_total function (counts_per_cell_after□=□10,000); the normalized counts were log-transformed using the pp.log1p function. The Pearson residuals were calculated from the raw counts for selecting top 5,000 highly variable genes using experimental.pp.highly_variable_genes function (n_top_genes=5,000). The StandardScaler function (with_mean=False) of scikit-learn v.0.24.0 was then used to scale the Pearson residuals of highly variable genes, followed by the TruncatedSVD function of scikit-learn v.0.24.0 to calculate the Principal Component Analysis (PCA) embeddings for the cells. The unsupervised graph-based Leiden clustering algorithm, pp.neighbors and tl.leiden functions, was used for the clustering based on the PCA embeddings. After clustering, COSG^64^ was used to identify top marker genes for each cluster, which were cross-compared with well-known marker genes from literature for cell type annotation. MAST^65^ was used to identify differentially expressed genes between different genotypes for each cell type. The R package clusterProfiler v4.2.1^66^ (pvalueCutoff = 0.01, qvalueCutoff = 0.01, and pAdjustMethod = “BH”) was then used to identify enriched ‘Biological Process’ Gene Ontology terms for the differentially expressed genes. GOplot v1.0.2^67^ with default parameters was used to calculate the z-scores of enriched Gene Ontology terms. TACCO^68^, with default parameters, was used to map the excitatory neuron identities between *Satb2 cko* and *Satb2 ctrl*, and the *Satb2 ctrl* snRNA-seq data was firstly filtered to retain only the union set of the top 100 marker genes per cell type, which was then used as the reference for TACCO mapping. For visualization of multiple snRNA-seq datasets across different conditions, we employed the piaso.pl.plot_embeddings_split from our single-cell analysis toolkit PIASO (https://github.com/genecell/PIASO) to align cell coordinates from different conditions and scale the gene expression or cell metrics for consistency.

### Heritability enrichment analysis with scDRS and LDSC

scDRS^69^ was used to identify intellectual deficiency (ID)– and intelligence-relevant cell types based on the scRNA-seq data with risk gene sets. Additionally, LDSC^23^ was used to identify ID-and intelligence-relevant cell types based on the scATAC-seq data with the genome-wide association study (GWAS) summary statistics. The ID gene list was downloaded from https://panelapp.genomicsengland.co.uk/panels/285/. For scDRS analysis, the GWAS MAGMA z-score weights were downloaded from https://figshare.com/articles/dataset/scDRS_data_release_030122/19312583, and top 1,000 genes with weights were used as the scDRS input. For LDSC analysis, we used liftOver with ‘-minMatch=0.5’ to lift mouse genomic regions (mm10) to human genome (hg19) to enable mapping the GWAS risk loci of human phenotypes. We then reciprocally converted the mapped regions (hg19) back to the mouse genome (mm10), retaining only those that mapped back to their original loci for LDSC analysis. The GWAS summary statistics and other input files required for LDSC analysis were downloaded from https://console.cloud.google.com/storage/browser/broad-alkesgroup-public-requester-pays/LDSCORE.

### Slide-seq V2 pre-processing and analysis

The sequenced reads of Slide-seq V2 libraries were aligned to GRCm39.103 reference and processed using the Slide-seq tools pipeline (https://github.com/MacoskoLab/slideseq-tools; v.0.2) to generate the gene count matrix and match the bead barcode between array and sequenced reads. We used RCTD^70^ (now available in the R package spacexr v2.2.1) to map cell types in Slide-seq V2 data based on reference snRNA-seq data. Only the genes detected in both the Slide-seq V2 data and the reference snRNA-seq data were retained in the reference dataset. The reference dataset was further filtered to retain only the union set of the top 100 marker genes per cell type identified by the R version of COSG (v0.9.0) with the following parameters: mu=100, remove_lowly_expressed=TRUE, expressed_pct=0.1. The accuracy of cell type decomposition in Slide-seq V2 data increased by the extraction of distinctive features for each cell type in the reference dataset.

### CTB labeling and AAV Tracing

Injections for tracing experiments were performed using either Alexa Fluor 555-conjugated cholera toxin subunit-B (CtB) (Invitrogen, C22843) at a concentration of 4 mg/mL in 1X PBS, or AAV9-FLEX-GFP (Salk Institute), injected through a pulled glass pipette attached to a Picospritzer III (Parker). For CTB experiments, littermate *Satb2^+/+^* and *Satb2^+/lacZ^*mice were used. For AAV experiments, *Satb2^+/lacZ^* mice were bred to various Cre-reporter mice: *Tlx3-Cre*, *Cux2-CreER*, and *Ntsr1-Cre*. Tamoxifen was administered to the *Cux2-CreER* mice at 150 mg/kg (dissolved in corn oil) daily from P25-P30 by oral gavage. Stereotaxic surgery was performed on P28 mice by anesthetizing them with isoflurane and placing them on a stereotaxic frame (Kopf). The skull was exposed, and coordinates were measured by using the bregma as the zero point. A craniotomy was performed, and the pulled glass pipette with CtB or AAV was placed on the brain surface to zero the Z coordinate. The needle was then lowered to the desired Z position, and 100 nL of CTB or 200 nL AAV was injected. After 5 minutes, the needle was retracted, and the skin was glued back together. Brains were collected and analyzed 5-days later for CTB and two weeks later for AAV. Injections were performed using the following coordinates: for WT: ventral posteromedial nucleus injection, AP –1.1 mm, ML 1.85 mm, Z –3.4 mm, cerebral peduncle injection AP –4 mm, ML 1.0 mm, Z –4.6 mm, corpus callosum injection AP –1.3 mm, ML 0.8 mm, Z –1.3 mm, SSp injection AP –1.3 mm, ML 3.3 mm, Z –0.9 mm. For Het: ventral posteromedial nucleus injection, AP –1.0 mm, ML 1.85 mm, Z –3.35 mm, cerebral peduncle injection AP –3.8 mm, ML 0.9 mm, Z –4.5 mm, corpus callosum injection AP –1.1 mm, ML 0.8 mm, Z –0.9 mm. For S1 injection AP –1.1 mm, ML 3.3 mm, Z –0.9 mm and CTB was continually injected as the needle was retracted from Z –0.9 mm to the Z –0.3 mm to deposit CTB across all cortical layers.

### Isolation induced ultrasonic vocalizations

Isolation induced ultrasonic vocalizations were recorded from P3, P7, and P14 mice using the Echo Meter Touch 2 (Wildlife Acoustics). Mouse pups were separated from their mother and quickly placed on fresh bedding in a new cage. The Echo Meter Touch 2 was positioned 10 cm above bedding and mice were recorded for 5 minutes before being returned to their home cage.

Ultrasonic vocalizations were analyzed with DeepSqueak^71^ and individual calls were labeled according to call categories identified by Fonseca, et al, 2021^47^. Call data was then imported into Prism for statistical analysis.

### Texture discrimination assay

On the first day, P30 mice were habituated in a chamber [40cm (L) by 40cm (W) by 40cm (H) open-field chamber] for 30 minutes. The next day, the mice were presented with two different textures for 5 minutes in the same chamber. Two 50 mL conical tubes were wrapped with 120-grit sandpaper (120-grit). The smooth tube (Sm) was then wrapped with cellophane to keep visual appearance of both tubes similar. Both tubes were weighed down with 50mL of water. Time spent interacting with each texture was measured when the nose of the mouse was within a 2cm proximity of each texture. Behavior was recorded using an overhead digital camera and tracked with Viewer III tracking system (BioServe).

Whisker trimming was performed as previously described^72^ and mystacial vibrissae were cut to skin level 24 hours before the texture preference test. Whisker trimmed mice were habituated and tested as stated above.

### Image acquisition and analysis

For cell counting analyses, high magnification images were acquired with a Zeiss 880 confocal microscope. Laser power and gain were adjusted until < 1% of pixels were saturated. Cell counting was performed on single z-slices. Images were divided into 500 μm wide regions and split into equally sized bins or cortical layers based on DAPI staining. An appropriate threshold was set for each channel and particles were analyzed with a circularity of 0.3-1.0 and size exclusion of > 1 μm. The same threshold was used across all images for the same channel between genotypes. Low magnification images were acquired using a Zeiss AxioImager Z2 widefield microscope. Statistical analysis was performed using GraphPad Prism 9.0. For each brain, the number of marker+ cells in the cortex were quantified in a 500 µm wide region from at least 3-4 matching sections per brain, from at least 3 brains per genotype. Care was taken to match the anterior-posterior, medial-lateral positions for the chosen areas between the mutant and control genotypes. Data are shown as mean ± SEM or ± SD as indicated in the figure legends. Statistical significance for single comparisons was determined using a nested t-test for comparisons of 3 or more conditions, or unpaired t-test for comparison of two conditions.

For fluorescence intensity measurements, images of whole sections were obtained using a Zeiss AxioImager Z2 widefield microscope with a 10X objective. The images were opened in QuPath, exported to imageJ (downsample 10), and saved as tif. The images were then opened in Fiji and the segmented line tool (width = 260 pixels) was used to select the contralateral cortex (non-injected side) from the midline to the entorhinal/piriform boarder. The straighten function was used to generate a rectangular representation of the selected region. The image was then split into 200 bins and mean signal intensity per bin was measured and normalized to the mean signal intensity of the injected area. The signal intensity across layers was calculated the same way, but in this case the selected area was 300 µm wide spanning the base of L6b to the pial surface. Fluorescence intensity means were determined by cortical layer (L1, L2/3, L4, L5a, L5b, L6), and the statistical significance of the difference of means was made via unpaired *t*-test using GaphPad Prism 9.0.

### Whole cell protein extraction

P0 cortices were dissected in ice cold 1X PBS with protease inhibitor (Sigma, 11873580001). The tissue was homogenized in standard RIPA buffer with protease inhibitors by manually triturating with a P1000 pipet tip before incubating on ice for 20 minutes. The cell homogenate was centrifuged at 14,000 g for 10 minutes at 4°C. The supernatant was removed and quantified via Pierce BCA Protein Assay Kit (Thermo Scientific, 23227). 10ug of protein was aliquoted and denatured at 100°C for 5 minutes in laemmli buffer (Bio-Rad, 1610747) supplemented with 2-Betamercaptoethanol.

### Co-Immunoprecipitations

P0 cortices were dissected in ice cold 1X PBS with protease inhibitor (Sigma, 11873580001). Nuclear extracts were isolated using the Active Motif Nuclear Complex Co-IP kit (Active Motif, 54001). 5 µg of the following antibodies used for precipitation of protein complexes; BCL11A (rabbit, Abcam, ab191401), BCL11B (rat, Abcam, ab18465), BRG1 (mouse, Santa Cruz Biotechnology, sc-17796), CHD4 (rabbit, Cell Signaling Technology 11912), CTCF (rabbit, Cell Signaling Technology 2899S), CUX1 (rabbit, Santa Cruz Biotechnology, sc-6327), EED (rabbit, Millipore, 09-774), HDAC1 (rabbit, Santa Cruz Biotechnology, sc-7872), HDAC2 (rabbit, Santa Cruz Biotechnology sc-7899), IGg (rabbit, Invitrogen, 02-6102), J1 (kindly provided by Jiang Wu, UTSW), LHX2 (goat, Santa Cruz Biotechnology, sc-19344), MBD2/3 (mouse, Santa Cruz Biotechnology sc-271562), MTA1 (rabbit, Cell Signaling Technology 5647), RBAP46 (rabbit, Cell Signaling Technology 6882), SATB2 (mouse, Abcam, Ab51502), SATB2 (rabbit, Abcam, ab34735). Antibody-protein complexes were precipitated with either Protein A or G magnetic beads (NEB, S1425S, S1430S) and eluted with a 0.1M Glycine, pH 2.5 buffer which was then neutralized with 1M Tris, pH 7.4. Samples were then denatured at 100°C for 5 minutes in laemmli buffer (Bio-Rad, 1610747) supplemented with 2-Betamercaptoethanol.

### Western blot analysis

The samples from whole cell extraction or co-IP were run on an 8% SDS-PAGE at 65V for 2.5 hours and transferred to a PVDF membrane (Sigma, IPVH85R) at 150 mA for 90 minutes. The membrane was blocked for one hour in 10 mL of 1% non-fat milk (Bio-Rad, 1706404) in 1X Tris-Buffered Saline, 0.1% Tween 20 (TBST). After blocking, 2 µg of primary antibody was added and incubated at 4°C overnight. The membrane was then washed 3x with TBST and incubated in Li-Cor IRDye 680RD or 800CW secondary antibodies at a 1:12,500 dilution for one hour. The membrane was washed 3x again with TBST, imaged on a Li-Cor imaging system, and images were processed using ImageStudioLite. Gapdh (mouse, Covance MMS-580S).

### Electrophysiology and neuronal morphology

Whole cell patch clamp recording was conducted in the SSp. To label the cortico-callosal-projecting (CC) SSp neurons, 50 nl of red retrobeads (Lumafluor) were injected into the contralateral S1 at least 24 h prior to recording. 350-μm thick coronal brain sections at SSp level were then made. Slices were cut in ice-cold ACSF (containing 126 mM NaCl, 2.5 mM KCl, 26 mM NaHCO_3_, 1 mM MgCl_2_, 2 mM CaCl_2_, 1.25 mM NaH_2_PO_4_, 10 mM glucose, and saturated with 95% O_2_ and 5% CO_2_). Slices were incubated at 34°C for 30 min and then transferred to the recording chamber. Bead^+^ CC neurons in layer 2/3 and layer 5 were visualized under a 60X objective (Olympus, NA = 0.9). Bead+ CC neurons with their soma at least 50 μm below the slice surface were selected to minimize truncation of dendritic arbors. The internal electrode solution contained: 130 mM K-gluconate, 10 mM HEPES, 4 mM KCl, 0.3 mM GTP-Na, 4 mM ATP-Mg, 2 mM NaCl,1 mM EGTA and 14 mM phosphocreatine (pH□7.2, 295-300□mOsm), with 0.15% (W/V) Neurobiotin (Vector Laboratories). To record miniature excitatory synaptic currents (mEPSCs), D-AP5 (50 μM, Tocris) and tetrodotoxin (TTX, 1μM, Tocris) were included in the circulating ACSF. Neuronal signals were conditioned using a MultiClamp 700B amplifier (Molecular Devices, Forster City, CA), low-pass filtered at 1 kHz (current) or 10 kHz (voltage signals), and digitized at 20 kHz using a Digidata 1440A digitizer under control of pClamp 10.6 software (Molecular Devices).

To reconstruct neuronal dendritic arbors, slices were fixed in 4% PFA overnight, extensively washed in PBS, and incubated with avidin-Alexa 488 (Invitrogen) for 24h in PBS containing 0.2% Triton X-100. Brain slices were then mounted on slides with a 350-um spacer to minimize crushing the tissue and better preserve Z information. Dendritic arbors were traced on an epifluorescence microscope (Zeiss M2) equipped with Neurolucida and a motorized stage (Prior Scientific, United Kingdom). Because apical dendrites were frequently cut off of L5 neurons, we only quantified dendritic arbors that are within 300um of the center soma. In comparison, full arbors of the L2/3 neurons were reported. Morphometric features were then extracted which included dendritic arbor, length, and number of intersections as a function of distances from soma using Neurolucida Explorer.

### Laser-scanning photostimulation and functional circuit mapping

Coronal slices at the level of SSp were obtained in *Satb2^+/+^*and *Satb2^+/lacZ^* littermate mice. Slices were perfused in modified ACSF (4 mM Ca^2+^, 4 mM Mg^2+^) containing 0.2 mM MNI-caged glutamate and 5 μM R-CPP to block NMDA receptors and potential short term plasticity changes. Bead+ cross projecting L5 or L2/3 CC neurons were then targeted for whole cell recording and LSPS mapping. Neurons with pyramidal-shaped soma were selected and voltage clamped at –70mV. Only neurons with a soma located at least 50 μm below the cut surface of the coronal slice were targeted for recording to minimize truncation of dendritic structure and connectivity.

LSPS mapping was performed with a 4x objective lens (NA 0.16; Olympus) after establishing whole cell recording using a 60x objective (NA 0.9; Olympus). Data acquisition and analysis were controlled by Ephus software^42^. 16X16 stimulation grids with 75 μm spacing were overlaid on the slice, with the top row of stimulation aligned with the pia surface. Digital images were registered using a CCD camera (Retiga 2000DC, Qimaging). 1-ms, 20 mW UV laser pulses was scanned onto slices unto the 16*16 stimulation locations. Laser power and timing were controlled by a mechanical shutter (Unibliz VCM-D1) and an optic shutter (Conoptics, model 3050). Prior to laser uncaging, neuronal firing properties were tested using a family of current step injections (–100 pA to 500 pA, with 50 pA increment). Membrane properties were constantly monitored with a –5 mV hyperpolarizing voltage step during uncaging. Neuronal signals were amplified with a Multiclamp 700B amplifier, digitized at 10 kHz, and acquired using two BNC-6259 boards (National Instruments, Austin, TX). Because L5 neurons receive major synaptic inputs to L2/3 neurons, in some experiments excitation profiles for L2/3 pyramidal neurons were tested^39,41^, during which loose-seal recordings were made from L2/3 neurons in current-clamp mode. The spike-generating sites for L2/3 neurons in response to glutamate uncaging (20mW, 1ms pulses) were mapped using an 8 × 8 stimulus grid with 50 μm spacing.

## Data Availability

ChIP-seq and CUT&RUN-seq data generated in this study have been deposited in the Gene Expression Omnibus (GEO) under the accession number GSE283035 and GSE320305. The snRNA-seq data for SSp cells from P28 *Satb2^+/fl^*(control), *Satb2^fl/lacZ^* (het), and *Satb2^fl/lacZ^; Emx1-Cre* (cKO) cortices have been deposited in GEO under the accession number GSE287066.

